# Outlier detection in multimodal MRI identifies rare individual phenotypes among 20,000 brains

**DOI:** 10.1101/2021.05.10.441017

**Authors:** Zhiwei Ma, Daniel S. Reich, Sarah Dembling, Jeff H. Duyn, Alan P. Koretsky

## Abstract

Outliers in neuroimaging represent spurious data or the data of unusual phenotypes that deserve special attention such as clinical follow-up. Outliers have usually been detected in a supervised or semi-supervised manner for labeled neuroimaging cohorts. There has been much less work using unsupervised outlier detection on large unlabeled cohorts like the UK Biobank brain imaging dataset. Given its large sample size, rare imaging phenotypes within this unique cohort are of interest, as they are often clinically relevant and could be informative for discovering new processes. Here we developed a two-level outlier detection and screening methodology to characterize individual outliers of multiple different brain imaging phenotypes from 20,000 UK Biobank subjects. In primary screening, every subject was parameterized with an outlier score per imaging phenotype to quantitate the degree of outlierness. This approach enabled the assessment of test-retest reliability via outlier scores, which ranged from excellent reliability for ventricular volume, white matter lesion volume, and fractional anisotropy, to good reliability for mean diffusivity and cortical thickness. Resting-state functional connectivity was eliminated for individual-level outlier screening due to its low test-retest reliability of outlier scores. In secondary screening, the extreme outliers (1110 subjects) were examined individually, and those arising from data collection/processing errors were eliminated. A subgroup (108 subjects) of the remaining non-artifactual outliers were radiologically reviewed, and radiological findings were identified in 98%. This study establishes an unsupervised framework for investigating rare individual imaging phenotypes within a large neuroimaging cohort.

## 1. Introduction

Outliers are defined as observations differing by a large amount from most other observations (Tan, Steinbach, & Kumar, 2006). By this definition, outliers constitute a small portion of a dataset and are exceptional patterns in some sense. Detecting outliers is of interest in brain imaging for two major reasons. First, outliers can occur due to imaging artifacts or noise. For example, head motion adversely affects brain morphometry, diffusion, and connectivity measurements (Power, Schlaggar, & Petersen, 2015; Reuter et al., 2015; Yendiki, Koldewyn, Kakunoori, Kanwisher, & Fischl, 2014) and causes outliers in these data. Second, and more importantly, some outliers represent unusual phenotypes that deserve special attention. For example, an anomalous MRI may indicate the presence of neurological disease that requires clinical attention. Certain unusual phenotypes may also be interesting for follow-up to determine the underlying mechanism for the large deviations of their brain MRI from the population.

Outlier detection methods applied in brain imaging can be categorized in many ways. One common way is based on whether the method makes use of labeled datasets to train the outlier detection model: supervised methods use labeled datasets that contain both labeled outliers and labeled non-outliers for training; semi-supervised methods use labeled datasets that only contain labeled non-outliers for training; and unsupervised methods use unlabeled datasets for training (Goldstein & Uchida, 2016). Using the available diagnostic labels for all subjects or at least the non-outlier subjects, outlier detection studies have employed a variety of algorithms, such as one-class support vector machine, Gaussian process regression, or autoencoders, and these have been applied in a supervised or semi-supervised manner to quantify the outlierness of healthy individuals or patients (Marquand, Rezek, Buitelaar, & Beckmann, 2016; Mourao-Miranda et al., 2011; Pinaya, Mechelli, & Sato, 2019; van Hespen et al., 2021). However, diagnostic labels are not always available, making the supervised or semi-supervised approaches challenging to implement across the board. Unsupervised outlier detection methods are needed for unlabeled brain imaging datasets, for example, the UK Biobank (UKB), an ongoing large epidemiological cohort (Miller et al., 2016).

The UKB is enrolling 500,000 subjects 40–69 years of age for extensive phenotyping and subsequent long-term monitoring of health outcomes (Allen et al., 2012). 100,000 subjects in this cohort are currently in the process of being invited back for MRI imaging, making it the largest multimodal MRI cohort in the world (Littlejohns et al., 2020). By enrolling the population of this age range, this unlabeled brain imaging dataset includes healthy and pre-symptomatic subjects, as well as a small fraction of subjects with different clinically relevant diseases. As time goes on, many more subjects in this cohort will become identified with a clinically relevant disease (Miller et al., 2016). Given its large sample size, the UKB cohort enables a unique opportunity for developing unsupervised outlier detection methods to discover rare imaging phenotypes. These rare imaging phenotypes could be clinically relevant or informative for discovering new processes and mechanisms.

In the present study, a two-level outlier detection and screening methodology was developed to characterize individual outlying MRI results across multiple brain imaging phenotypes among 20,000 UKB subjects. We made use of the multimodal MRI data to derive ventricular, white matter, and gray matter-based imaging phenotypes of the brain (Fig. 1a). Every subject was parameterized with an “outlier score” per imaging phenotype in an unsupervised manner without any prior labels (Fig. 1b). This outlier score quantifies how far an individual deviates from most other subjects. Test-retest reliability of outlier scores of each imaging phenotype was characterized in the subjects that had repeat MRI scans, and any less reliable imaging phenotype was not used for further individual-level outlier screening. Individual extreme outlier subjects were categorized according to whether there were data collection/processing errors, or whether the individual had radiological findings or appeared normal as determined by a board-certified neuroradiologist (Fig. 1c). Similar outlier detection and screening procedures were also carried out separately in the Human Connectome Project (HCP) dataset, and the extreme outlier subjects from this young adult cohort that might be interesting for follow-up are also described.

**Fig. 1.**
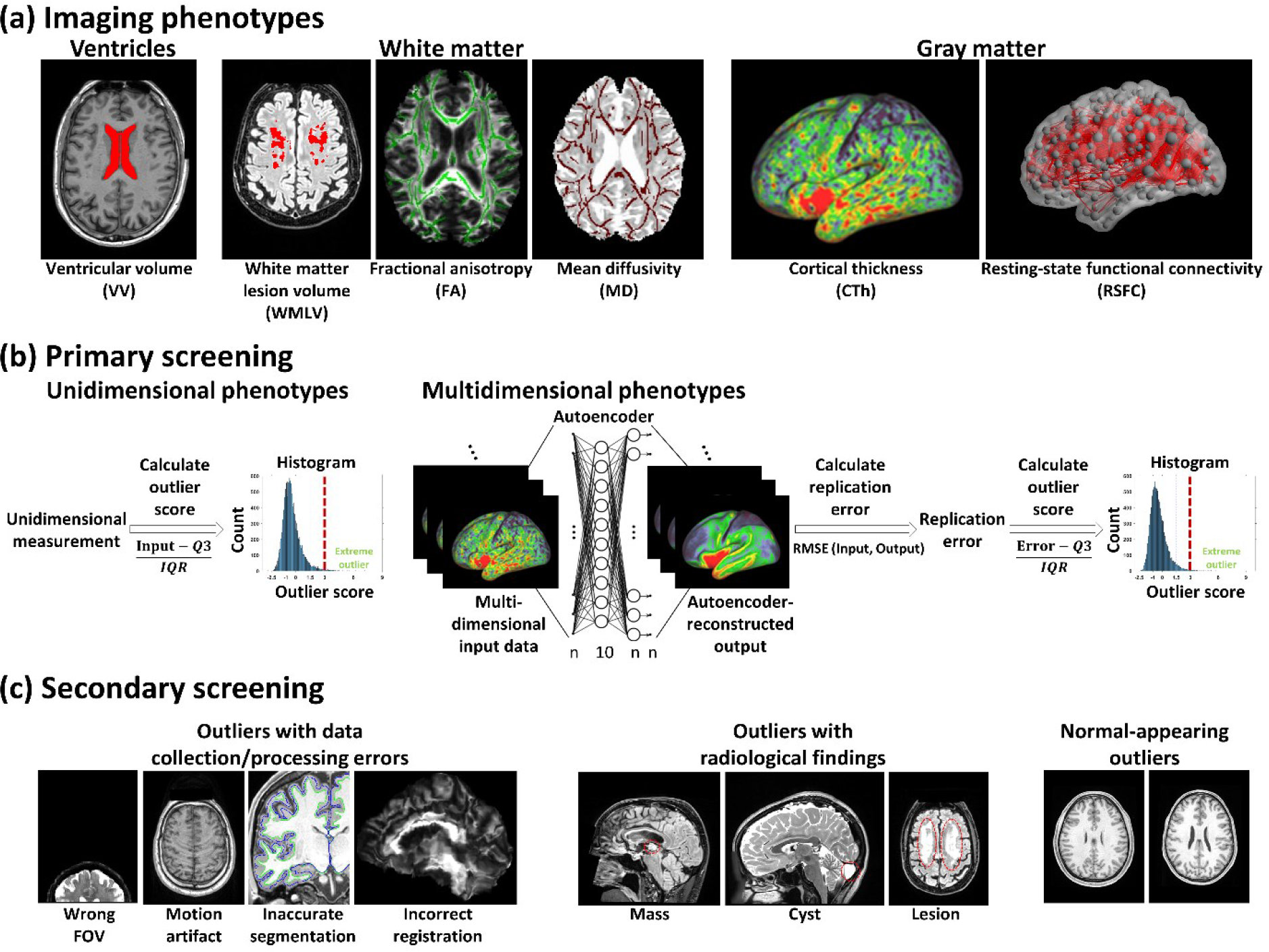
Schematic illustration of outlier detection and screening pipeline. (**a**) Brain imaging phenotypes used for outlier detection. (**b**) Primary screening: calculation of outlier scores. (**c**) Secondary screening: investigation of individual extreme outliers.

## 2. Materials and Methods

### 2.1 Main dataset

The brain imaging data of 19411 UKB subjects (9172 males and 10239 females; age 44–80) at the initial imaging visit were used in the present study (hereafter referred to as *UKB discovery group*). If available (not marked as “unusable” or “incompatible” by the UKB), each subject’s T1w MPRAGE and T2w FLAIR structural MRI, spin echo (SE) echo-planar imaging (EPI) diffusion MRI (dMRI), and gradient echo (GE) EPI resting-state functional MRI (rsfMRI) data were used. Some subjects only had usable structural MRI data, resulting in a reduced sample size of dMRI and rsfMRI data. The detailed demographic information is summarized in Table S1. The data were acquired on identical 3T Siemens Skyra MRI scanners, and the detailed acquisition protocols can be found elsewhere (Alfaro-Almagro et al., 2018). The UKB study was approved by the North West Multi-centre Research Ethics Committee, and informed consent was obtained from all participants. The present study was approved by the Office of Human Subjects Research Protections at the National Institutes of Health (ID#: 18-NINDS-00353).

**Table 1.** Summary of outlier score distributions for the main dataset (UKB discovery group).

| Phenotype | VV | WMLV | FA | MD | CTh | RSFC* |
| --- | --- | --- | --- | --- | --- | --- |
| Number of subjects | 19411 | 18467 | 17947 | 17947 | 18393 | 17220 |
| Skewness | 1.77 | 4.36 | 7.54 | 9.04 | 1.69 | 1.51 |
| Kurtosis | 10.06 | 37.27 | 191.12 | 178.31 | 9.54 | 12.01 |
| Number of outliers | 158<br>(0.8%) | 716<br>(3.9%) | 189<br>(1.1%) | 258<br>(1.4%) | 119<br>(0.6%) | 60<br>(0.3%) |

\*Note: full correlations without global signal regression.

### 2.2 Image preprocessing and extraction of imaging phenotypes

The following six commonly used brain imaging phenotypes were extracted from imaging preprocessing outputs: ventricular volume (VV), white matter lesion volume (WMLV), fractional anisotropy (FA), mean diffusivity (MD), cortical thickness (CTh), and resting-state functional connectivity (RSFC). The detailed procedures are described as follows.

The raw T1w MPRAGE and T2-FLAIR images were preprocessed by the HCP structural pipeline (v4) (Glasser et al., 2013) based on FreeSurfer (v6) (Fischl, 2012). For the subjects without usable T2-FLAIR images, the ventricles were segmented from their T1w images using FreeSurfer. The ventricular segmentations were manually inspected for each subject, and the ones with large segmentation defects in their enlarged ventricles were reprocessed with “-bigventricles” flag in FreeSurfer to correct the defects. Each subject’s VV was calculated by summing the volumes of lateral ventricles, temporal horns of the lateral ventricles, choroid plexuses, third ventricle, and fourth ventricle. WMLV was calculated by the Brain Intensity Abnormality Classification Algorithm (BIANCA) (Griffanti et al., 2016), a k-nearest-neighbor-based automated supervised method, using T2-FLAIR images but also T1w images as its inputs. CTh values in the standard CIFTI grayordinate space (with folding-related effects corrected) from only the subjects preprocessed successfully by the HCP structural pipeline, were used for primary screening.

The dMRI data underwent FSL eddy-current and head-movement correction (Andersson & Sotiropoulos, 2016), gradient distortion correction, diffusion tensor model fitting using the b = 1000 shell (Basser, Mattiello, & LeBihan, 1994), and Tract-Based Spatial Statistics (TBSS) analyses (Smith et al., 2006). The TBSS skeletonized images were averaged within the ROI of the John Hopkins University white matter atlas (Mori et al., 2008). Here the original MD values were multiplied by 10000 to convert to the unit of 10^-4^ mm^2^/s. The FA or MD maps of 27 major white matter ROIs (Table S2) were used for primary screening.

**Table 2.**
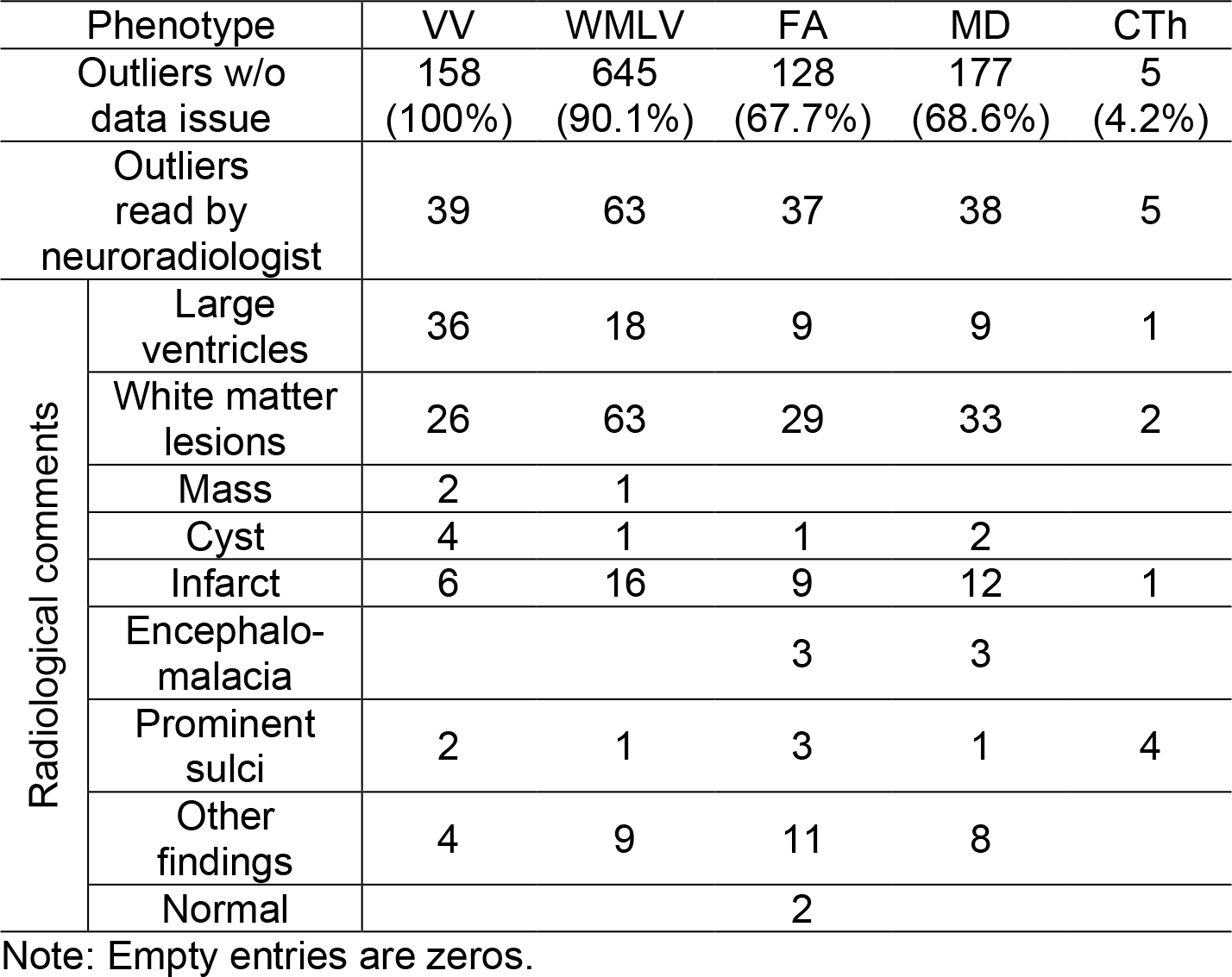
Summary of radiological review results of the outlier subjects in the main dataset.

The rsfMRI data were preprocessed by the UKB rsfMRI pipeline (v1) (Alfaro-Almagro et al., 2018), and the volumetric FIX-denoised data (Griffanti et al., 2014; Salimi-Khorshidi et al., 2014) were brought to the standard CIFTI grayordinate space using Ciftify (v2.3.2) (Dickie et al., 2019). For each subject, the standard deviation (SD) of percent change time series of each grayordinate was calculated, and the grayordinates with this SD greater than 0.1 were considered as noisy grayordinates. These noisy grayordinates were masked from further analyses. Using a well-established RSFC-based parcellation scheme (333 parcels) (Gordon et al., 2016), RSFC was quantified by the Pearson cross-correlation coefficient between the ROI-averaged time series of each pair of parcels, with or without global signal regression, respectively. In addition, RSFC was quantified using partial correlations with Tikhonov regularization (ρ = 0.5; FSLNets) (Pervaiz, Vidaurre, Woolrich, & Smith, 2020). Due to the symmetry of the RSFC matrices, the upper triangular parts of these matrices (333*332/2 = 55278 elements) from each of these three RSFC evaluation methods were used for primary screening respectively.

### 2.3 Primary screening: calculation of outlier scores

In primary screening, every subject was parameterized with an outlier score per imaging phenotype. The outlier score quantified the degree of outlierness in that imaging phenotype, and extreme outliers were identified based on the outlier scores. In statistics, extreme outliers in distribution are defined as the observations above the third quartile (*Q3*) plus three times the interquartile range (*IQR*) of that distribution (Tukey, 1977). For a unidimensional imaging phenotype (VV, WMLV), using VV as an example, the number of *IQR*s away from the *Q3* of the VV distribution was used to define VV outlier scores:

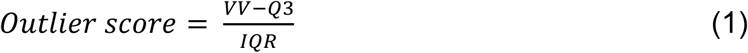

In this way, the unit of outlier score is *IQR*, and an extreme outlier has an outlier score of greater than 3. WMLV outlier scores were calculated similarly.

For each multidimensional imaging phenotype (FA, MD, CTh, RSFC), an autoencoder was used to calculate the outlier scores (Hawkins, He, Williams, & Baxter, 2002). Setting the dimensionality of the imaging phenotype as M and the number of subjects in the UKB discovery group as N, the inputs to the autoencoder were the values of that imaging phenotype across the whole group (M * N), and the autoencoder was trained to replicate this input at its output. By definition, outliers only comprised a small portion of a dataset, therefore the trained autoencoder cannot replicate these outliers as good as the non-outliers. This resulted in larger replication errors for the outlying subjects. These replication errors (also known as “autoencoder reconstruction error”) were measured by the root mean square errors between each input and the autoencoder-predicted output. Because these replication errors were unidimensional, similar to the calculation of outlier scores for unidimensional imaging phenotypes, the number of *IQR*s away from the *Q3* of the replication error distribution was used to define outlier scores:

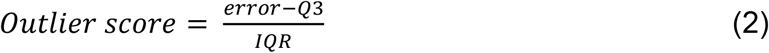

Still, the unit of outlier score is *IQR*, and an extreme outlier has an outlier score of greater than 3.

In the above analyses, to control for the effects of two covariates (age, brain volume) on outlier detection, their correlations with VV, WMLV, and the autoencoder replication errors of multidimensional imaging phenotypes were evaluated. The covariates with correlation >0.3 were regressed out from VV, WMLV, or the replication errors before applying Eq. (1) or (2). As a result, brain volume and age were regressed out from VV, and only age was regressed out from WMLV (Fig. S1).

Each autoencoder used in the present study was comprised of an input layer (M dimensions), a hidden layer of 10 neurons, and an output layer (M dimensions). A sparsity proportion of 0.05 was used, and the sparsity regularization coefficient was set to 1. The L2 weight regularization coefficient was set to 0.001. The sigmoid function was used as the activation function, and the mean squared error function adjusted for sparse autoencoder was used as the loss function. A scaled conjugate gradient descent algorithm (Moller, 1993) was used for training the autoencoder. The autoencoders were implemented using the ‘trainAutoencoder’ function in the MATLAB and were trained using a GPU cluster (https://hpc.nih.gov). When the input dataset was too large to fit into the GPU memory, multiple autoencoders were used: In these scenarios, the input data were split into 9 to 10 smaller subgroups in a stratified manner, preserving the ratio of age and sex in each subgroup. For each subgroup, an autoencoder was trained using the data of that subgroup as the input. The trained autoencoders were then applied to the full dataset and the output of the whole group was obtained by averaging the outputs from each of these autoencoders.

### 2.4 Evaluation of reliability of outlier scores and elimination of less reliable imaging phenotype

A subgroup (1427 subjects) of the UKB discovery group subjects had a repeat MRI session (aka “retest”) two to three years after the initial imaging visit (aka “test”). The test and retest data of these subjects were used to evaluate long-term reliability of outlier scores. For each unidimensional imaging phenotype, unlike the primary screening, their measurements in the reliability analysis were no longer adjusted for covariates. The *Q3* and *IQR* were calculated from the full test data and applied to calculate outlier scores for both test data and, for subjects who were scanned twice, retest data. For each multidimensional imaging phenotype, outlier scores of the test visit calculated in the primary screening were used directly. To calculate the outlier scores of the retest visit, the autoencoders trained on the full test data were applied to the retest data. The reliability was quantified by intraclass correlation coefficient (Shrout & Fleiss, 1979) (ICC) between the outlier scores of the test and retest data using a one-way random effects model:

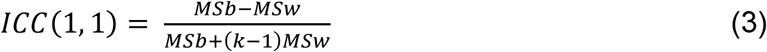

where *MSb* is the between-subject mean square, *MSw* is the within-subject mean square, and *k* is the number of observations per subject (McGraw & Wong, 1996). Reliability was defined as excellent (ICC > 0.8), good (0.8 > ICC > 0.6), moderate (0.6 > ICC > 0.4), fair (0.4 > ICC > 0.2), or poor (ICC < 0.2) (Guo et al., 2012) in the present study.

Any imaging phenotype with moderate/fair/poor outlier score reliability was excluded from further analysis of individual outliers. This resulted in the exclusion of RSFC (for details, see *3.2 Long-term test-retest reliability of outlier scores*).

### 2.5 Secondary screening: investigation of individual extreme outliers

For each remaining imaging phenotype, the extreme outlier subjects were first checked to see if they were associated with data collection/processing errors.

For each VV extreme outlier subject, ventricle segmentation quality was visually inspected by overlaying the border of the segmented ventricle mask on the T1w image. For each WMLV extreme outlier subject, white matter lesion segmentation quality was visually inspected by overlaying the border of the segmented lesion mask on the T2-FLAIR image. The VV or WMLV extreme outlier subjects with incorrect segmentation were deemed to be associated with data collection/processing errors.

For the subjects with usable dMRI data, the dMRI motion parameters (*.eddy_restricted_movement_rms) were calculated by FSL’s eddy tool (Andersson & Sotiropoulos, 2016), and the head motion of each subject was summarized by the mean and largest values of the volumetric movements between adjacent frames. The subjects with at least one of these two summary parameters above the upper inner fence (*Q3* + 1.5 * *IQR*, commonly used to define mild outliers in statistics (Tukey, 1977)) of the distribution were flagged as having severe head motion. The registration quality was assessed by each subject’s mean deformation of the TBSS nonlinear registration, and the subjects with this parameter above the upper inner fence (*Q3* + 1.5 * *IQR*) of the distribution were flagged as having bad registration. The FA or MD extreme outlier subjects were also visually checked for registration quality and FOV coverage. Taken together, the FA or MD extreme outlier subjects flagged as having incorrect FOV coverage, severe head motion, or bad registration were deemed to be associated with data collection/processing errors.

For CTh, volume registration quality was quantified by the number of suprathreshold voxels in the Jacobian map of nonlinear registration. Surface registration quality was quantified by areal and shape distortion maps of folding alignment (MSMSulc) surface registration (Robinson et al., 2018). Using the aforementioned multidimensional outlier detection method and these distortion maps as inputs, an outlier score for the areal distortion map and an outlier score for the shape distortion map were calculated for each subject. T1w/T2w ratio myelin maps (Glasser & Van Essen, 2011) were further used to detect potential surface segmentation issues that could be caused by the subject’s anatomy, and an outlier score for the T1w/T2w ratio myelin map was calculated for each subject via multidimensional outlier detection. White/pial surface segmentation quality of CTh extreme outlier subjects was checked via HCP pipeline structural quality control scenes (https://github.com/Washington-University/StructuralQC; v1.4.0), and the CTh extreme outlier subjects with poor surface segmentation were flagged. T1w structural images of the CTh extreme outlier subjects were also inspected visually, and the subjects with visible motion artifacts such as ringing artifacts were flagged. Taken together, the CTh extreme outlier subjects with bad volume or surface registration quality, anomalous T1w/T2w ratio map, bad white/pial surface segmentation, or visible motion artifacts in T1w images were deemed to be associated with data collection/processing errors.

A subgroup (108 subjects) of the remaining extreme outlier subjects without data collection/processing errors were radiologically reviewed. The subjects in this subgroup were manually sampled to not only include all top-ranked outlier subjects but also ensure a wide coverage of non-top extreme outlier subjects in each imaging phenotype (Fig. S2). T1w MPRAGE and T2-FLAIR images, as well as the ages of these subjects, were provided to a board-certified neuroradiologist (D.S.R.). The instructions to the neuroradiologist were to identify major findings that might plausibly account for the extreme outlier score — not to identify subtle abnormalities that would have required dedicated review on clinical-grade display systems. When the neuroradiologist had an uncertain diagnosis, UKB health outcomes data (UKB Category 1712) were used in an attempt to determine the diagnosis. This data recorded the first occurrence of various diseases, including neuropsychiatric and neurological disorders. Based on the radiological review results, the subjects in the subgroup were further divided into two subgroups: a subgroup of the extreme outlier subjects with radiological findings (106 subjects), and another subgroup of the extreme outlier subjects appeared normal to the neuroradiologist (2 subjects). The cases from these two subgroups that would be interesting for follow-up were highlighted.

### 2.6 Evaluation of the relationships between outlier scores of different imaging phenotypes

The relationships between outlier scores of different imaging phenotypes were quantified using Pearson cross-correlation coefficients in the UKB discovery group. The extreme outliers due to data collection/processing errors were excluded from this analysis. Two representative relationships of outlier scores, WMLV versus VV, and WMLV versus FA, were also visualized using scatterplots. In each scatterplot, three zones were defined to categorize extreme outlier subjects. For WMLV versus VV, zone I covered the subjects who were VV extreme outliers but with normal WMLV (WMLV outlier score < 1.5), zone II covered the subjects who were both VV and WMLV extreme outliers, and zone III covered the subjects who were WMLV extreme outliers but with normal VV (VV outlier score < 1.5). The density of subjects in each zone was calculated by dividing the number of subjects by the area of the zone as follows:

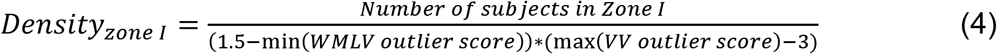

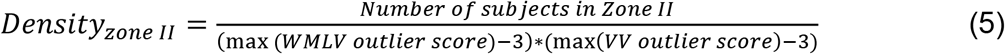

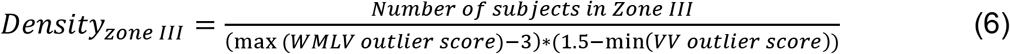

To evaluate the differences in densities across the three zones, a bootstrap procedure with replacement on subjects was used to generate 100,000 bootstrap samples of the original sample size. For each bootstrap sample, the density of each zone was re-computed. A one-way ANOVA was then performed to evaluate the differences across the zones using the bootstrap samples. Similar analyses were also carried out to evaluate the relationship between WMLV and FA outlier scores.

### 2.7 Outlier detection and screening in the HCP dataset

Similar outlier detection procedures were carried out separately in the HCP dataset to identify interesting extreme outliers in this young adult cohort (for details, see *Supplementary Text*). Briefly, 3T MRI data from the 1200 Subjects Release (1113 subjects: 550 males and 656 females; age 22–37) were used. Because of the lack of HCP T2-FLAIR data and poor WMLV segmentation accuracy when only using T1w images (Hotz et al., 2021), WMLV was excluded from the outlier detection of the HCP dataset. All the HCP extreme outliers (11 subjects) without data collection/processing errors were radiologically reviewed, and the cases that would be interesting for follow-up were highlighted.

## 3. Results

### 3.1 Properties of outlier score distributions

The results presented throughout the rest of the manuscript were obtained using the UKB discovery group unless otherwise specified. The outlier score histogram of each imaging phenotype is shown in Fig. 2. These distributions were all right-skewed and more leptokurtic than a standard normal distribution (see Table 1 for skewness and kurtosis values). The percentage of extreme outliers ranged from a lowest of 0.3% in RSFC, to a highest of 3.9% in WMLV (Table 1). These percentages are all much higher than a standard normal distribution predicts, because the criterion of *Q3* + 3 * *IQR* for defining extreme outliers (referred to as “outlier” hereafter) in each distribution is equivalent to about 4.7 times the SD plus the mean in a standard normal distribution. One would predict only 0.0001% of the data above *mean + 4.7 * SD* in a standard normal distribution. Taken together, the results suggest that the outlier score distributions were all more outlier-prone than a standard normal distribution.

**Fig. 2.**
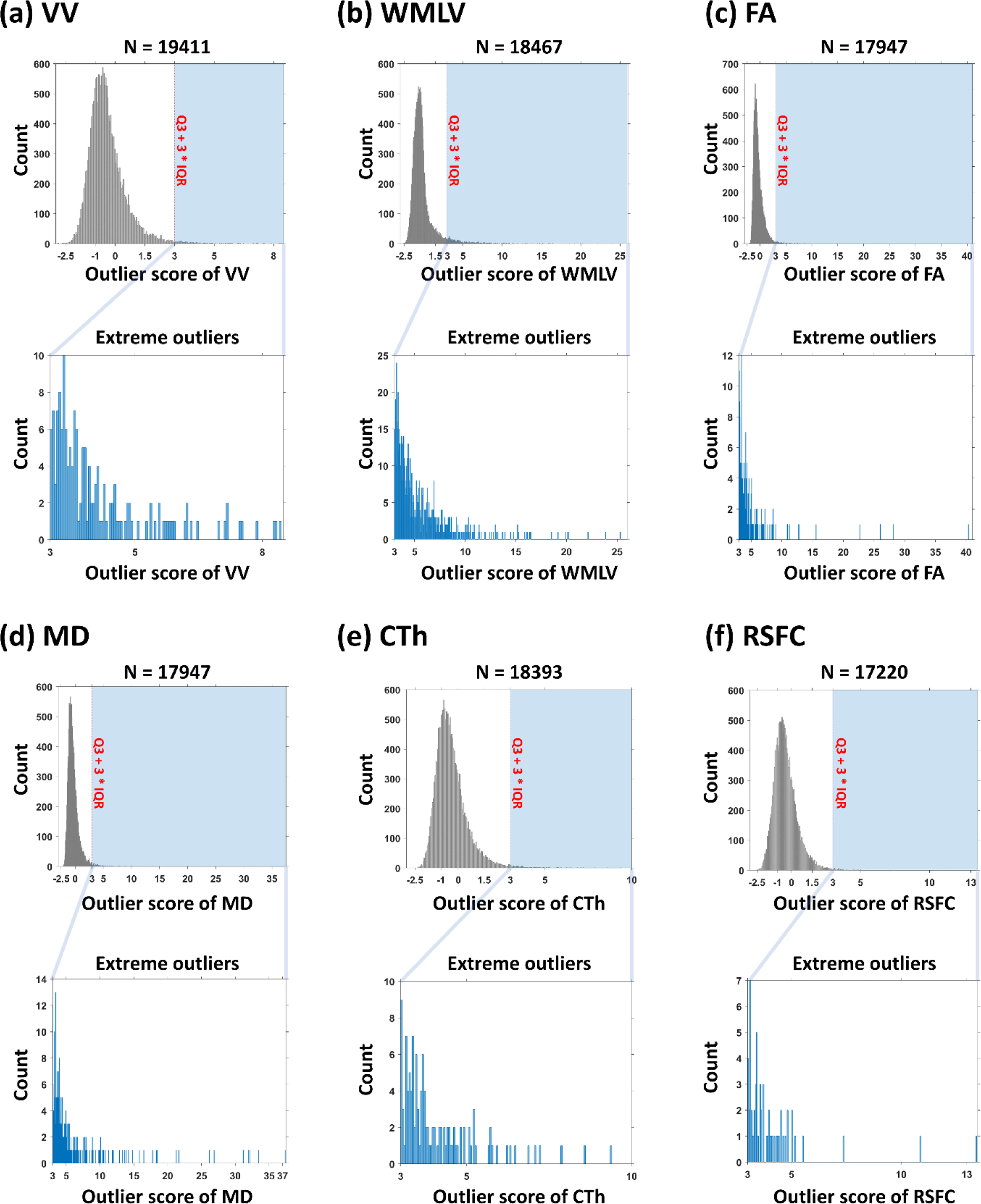
Outlier score histograms. (**a**) VV. (**b**) WMLV. (**c**) FA. (**d**) MD. (**e**) CTh. (**f**) RSFC. The zoom panels show the outlier score histograms of extreme outlier subjects.

### 3.2 Long-term test-retest reliability of outlier scores

A subgroup of the discovery group subjects had a repeat MRI session two to three years after the initial visit. The outlier scores of test versus retest of each imaging phenotype are visualized in the scatterplots of Fig. 3a-f, respectively. VV outlier scores had excellent test-retest reliability, as indicated by the close-to-one value of the ICC (ICC = 0.98) between test and retest outlier scores. The test-retest reliabilities of WMLV and FA outlier scores were lower than VV but still excellent (WMLV ICC = 0.82; FA ICC = 0.87). The test-retest reliabilities of MD and CTh outlier scores were lower than the former three but still in the range of good reliability (MD ICC = 0.75; CTh ICC = 0.62).

**Fig. 3.**
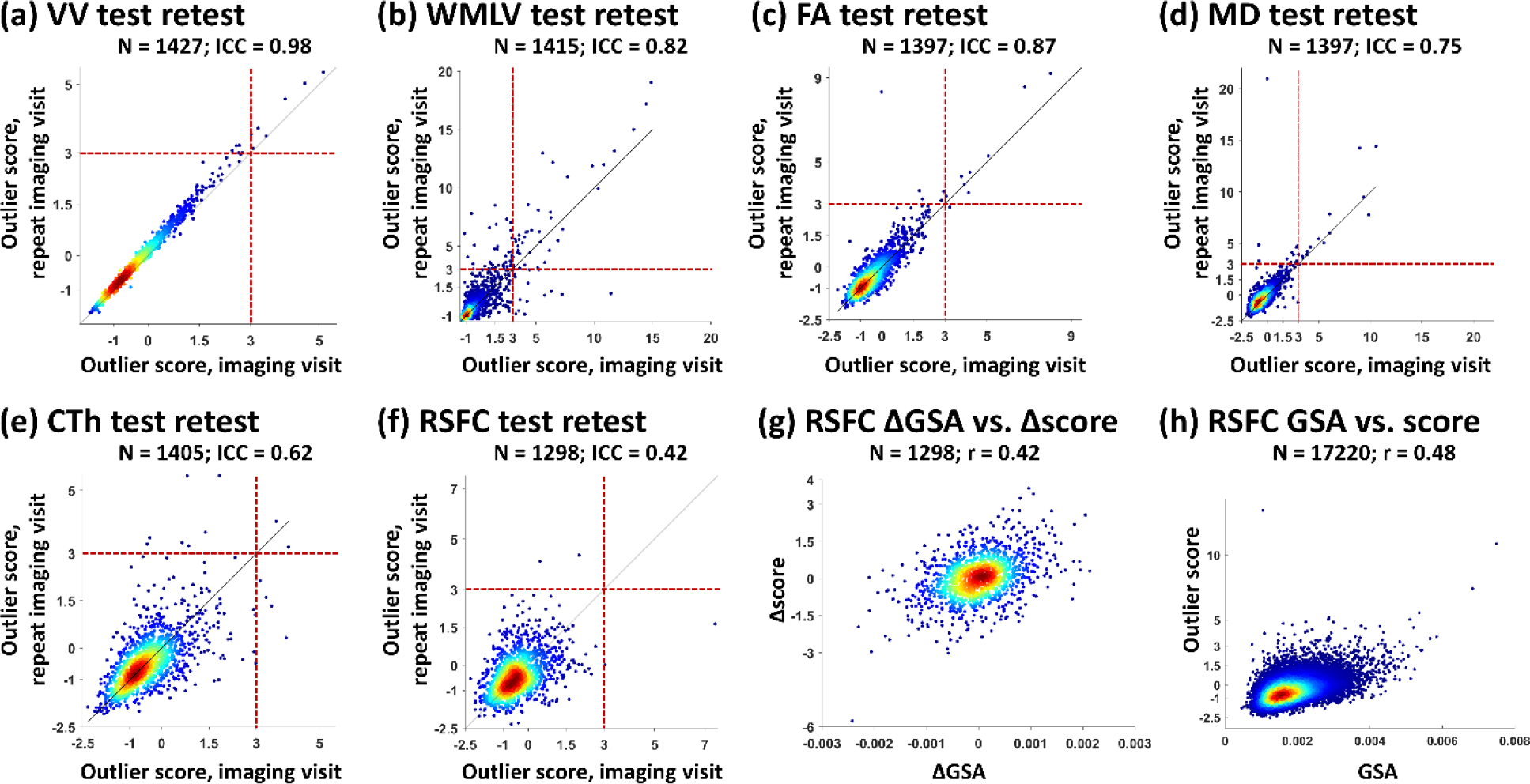
Long-term test-retest reliability of outlier scores. (**a**) VV. (**b**) WMLV. (**c**) FA. (**d**) MD. (**e**) CTh. (**f**) RSFC. For **a**-**f**, in each scatterplot, each subject’s outlier score of the initial imaging visit (aka “test”; year 2014+) is plotted against the outlier score of the first repeat imaging visit (aka “retest”; year 2019+). ICC: intraclass correlation between outlier scores of the two visits. Red dashed line: *Q3* + 3 * *IQR*. (**g**) The scatterplot of test-retest global signal amplitude (GSA) change versus test-retest RSFC outlier score change. For (a)-(g), only the UKB subjects that had both test and retest data available are shown in these scatterplots. (**h**) The scatterplot of GSA versus RSFC outlier score (RSFC calculated using full correlations).

However, RSFC outlier scores had a low test-retest ICC (ICC = 0.42. Fig. 3f). Because of this low reliability, among the subjects with available test-retest data, no subject had both test and retest RSFC identified consistently as an outlier. This change in test-retest outlier scores was found to be correlated with the change of global signal amplitude (r = 0.42. Fig. 3g). Here, global signal amplitude was defined as the SD of the global signal (Wong, Olafsson, Tal, & Liu, 2013). Indeed, the RSFC outlier score itself was found to be moderately correlated with global signal amplitude (r = 0.48. Fig. 3h). This association was not due to head motion, because such correlation persisted after excluding subjects with large head motion (r = 0.51. Fig. S3a). The association between RSFC outlier score and global signal amplitude also persisted when using partial correlations to evaluate RSFC, although they became negatively correlated in this case (r = -0.70. Fig. S3b). Global signal regression reduced their association, but RSFC outlier score was still weakly correlated with global signal amplitude (r = 0.39. Fig. S3c). Remarkably, when we carried out similar analyses on the HCP dataset, the results were very similar (Fig. S3d-i). Thus, RSFC was eliminated for further individual-level outlier screening due to its low individual test-retest reliability.

### 3.3 Summary of the screening results of individual outliers

The total number of outliers across all individual imaging phenotypes (excluding RSFC) was 1440. Because there were subjects who were outliers in more than one imaging phenotype, there were 1110 distinct subjects that made up these 1440 outliers.

Through the screening of each outlier subject, 211 of these distinct subjects were associated with data collection/processing errors. Interestingly, none of the VV outliers were associated with data collection/processing errors. More frequent data collection/processing errors were found in the WMLV, FA, or MD outliers as compared to the VV outliers (Table 2). Some of these errors occurred at the data acquisition stage, due to head motion artifacts (Figs. S4a and S5b) or the selection of a wrong FOV (Fig. S5a). Others occurred at the data processing stage, such as incorrect segmentation (Fig. S4b) or incorrect registration (Fig. S5c). Data collection/processing errors were found to be most common in the CTh outlier subjects. Indeed, 95.8% (114/119) of the CTh outlier subjects had data collection/processing errors. Similar to the errors found in the white matter outliers, these errors were due to head motion during data collection (Fig. S6a), incorrect segmentation/registration in data processing (Figs. S6b and S6c), or the combination of these issues (Fig. S6d).

Of the remaining 899 subjects that did not have data collection/processing errors, 108 were reviewed by a board-certified neuroradiologist (Table 2). These 108 subjects included all top-ranked outlier subjects and a diverse number of non-top-ranked outlier subjects (see Fig. S2 for details), which spanned almost the whole range above *Q3* + 3 * *IQR* of the outlier score distribution in each imaging phenotype. In this subgroup, 106 subjects (98.1%, 106/108) were identified with radiological findings, and these findings covered a diverse category of phenotypes, such as large ventricles, masses, cysts, white matter lesions, infarcts, encephalomalacia, and prominent sulci. Representative individual outlier subjects are reported in the next few subsections per imaging phenotype.

### 3.4 Individual outliers of ventricular volume

As an example, Fig. 4a shows a VV outlier subject versus a normal subject. This subject had significantly enlarged lateral ventricles compared to a normal one (about 8.2 * *IQR* away between these two subjects in VV outlier score distribution). Thirty-nine of the VV outliers were reviewed by the neuroradiologist. All the VV outliers being read were identified with radiological findings of large ventricles. Some of them had relatively clear etiology: A third ventricle mass (possibly a choroid plexus papilloma), a fourth ventricle mass (possibly an ependymoma), a colloid cyst, and a frontoparietal arachnoid cyst, all of which could cause obstructive hydrocephalus, were found in four VV outlier subjects respectively (Fig. 4b). The other major pathologies identified in the VV outliers were infarcts, nodules, agenesis of corpus callosum, and white matter lesions (Fig. S7a).

**Fig. 4.**
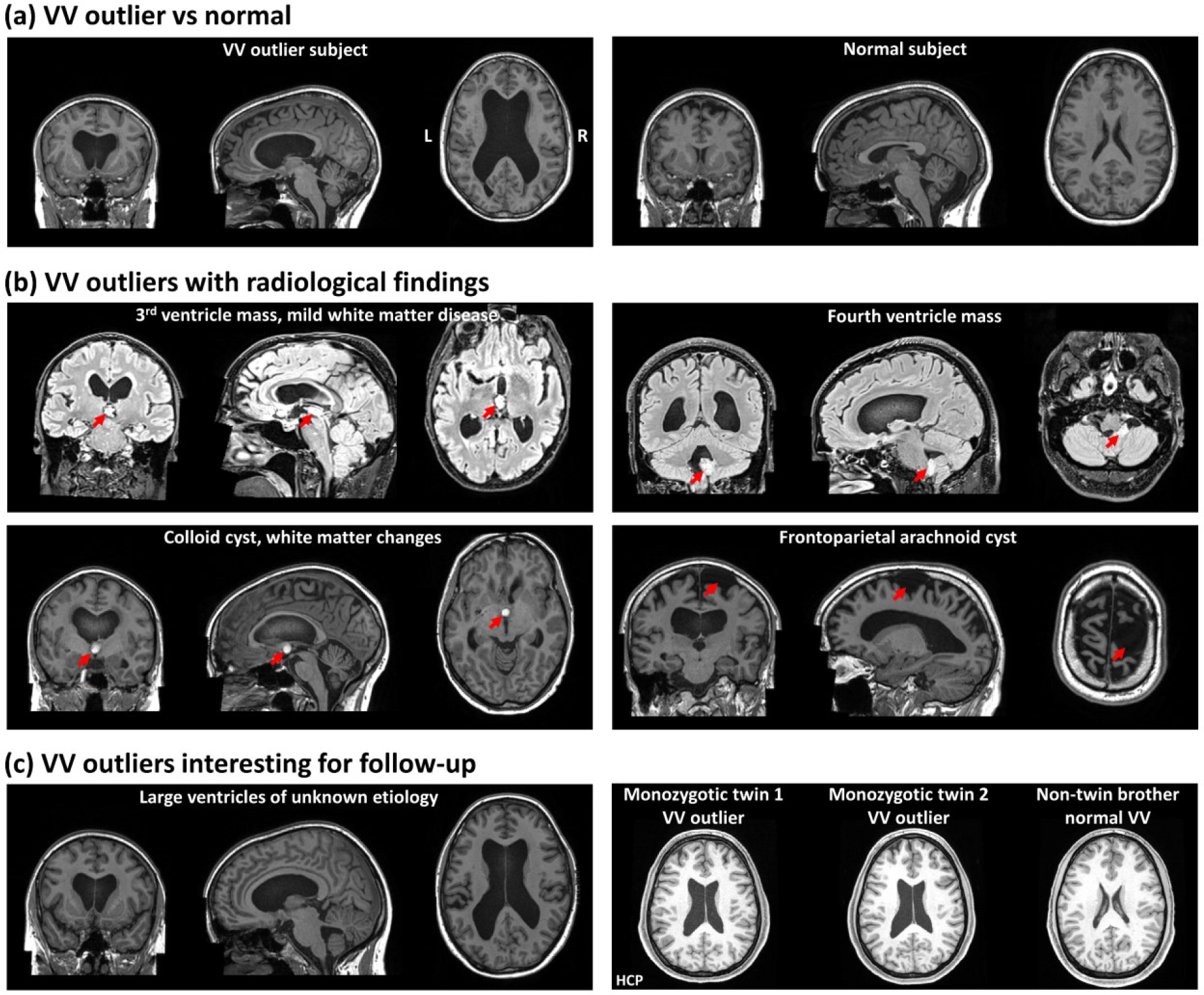
Individual outliers of VV. (**a**) Structural images of an example of a VV outlier subject (left column) and an example of a normal VV subject (right column). (**b**) Structural images showing radiological findings in four representative VV outlier subjects. (**c**) Structural images of VV outlier subjects interesting for follow-up. Left column: a subject with large ventricles of unknown etiology. Right column: Structural images of a family in the HCP dataset (monozygotic twins and their non-twin brother). The twins had large ventricles of unknown etiology, but their non-twin brother had normal VV.

In addition, a few VV outliers being read would be potentially interesting for follow-up because they had large ventricles of unknown etiology and they did not present any other noticeable pathology (Fig. 4c, left panel). Such VV outliers of unknown etiology were also present in the HCP dataset. In one family (Fig. 4c, right panel), female monozygotic twins were both VV outliers, but their non-twin brother had normal VV; in another family (Fig. S7b), one twin of a male monozygotic twin pair was a VV outlier, but the other twin and his non-twin brother both had normal VV. These twin data open the possibility of probing genetic and environmental causes underlying the anomalously large VV. Taken together, the results indicate VV outliers were associated with multiple different brain pathologies, and some of them had uncertain etiology requiring additional follow-up investigation.

### 3.5 Individual outliers of white matter-based imaging phenotypes

Outlier detection of white matter-based imaging phenotypes was performed with WMLV, FA, and MD, respectively. As an example, Fig. 5a shows a WMLV outlier subject versus a normal subject (about 26.5 * *IQR* away between these two subjects in WMLV outlier score distribution). The outlier subject had irregular periventricular white matter lesions extending into the deep white matter with large confluent areas, whereas the example normal subject had only tiny lesions on the periventricular caps. Fig. 5b shows regional FA deviation maps of an FA outlier subject versus a normal subject (about 9.5 * *IQR* away between these two subjects in FA outlier score distribution). For this representative outlier subject, regional FA negatively deviated in all 27 white matter ROI used in this study, whereas the FA of the representative normal subject had almost no deviations. Fig. S8a shows regional MD deviation maps of an MD outlier subject versus a normal subject (about 5.5 * *IQR* away between these two subjects in MD outlier score distribution), in which a large positive MD deviation was observed in the left superior longitudinal fasciculus of this outlier subject.

**Fig. 5.**
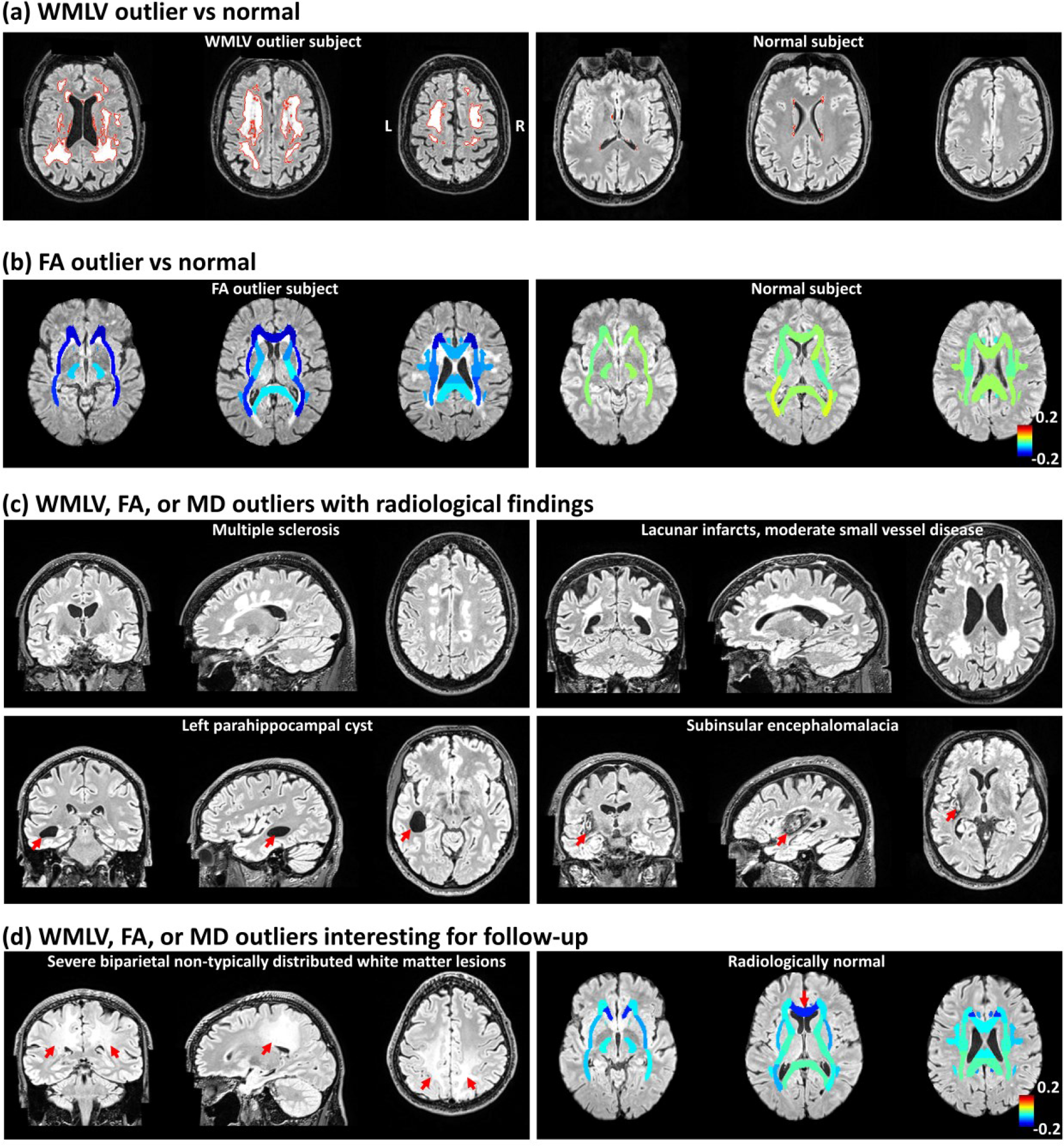
Individual outliers of white matter-based imaging phenotypes. (**a**) T2 FLAIR images of an example of a WMLV outlier subject (left column) and an example of a normal WMLV subject (right column). The red line represents the boundary of white matter lesions. (**b**) Regional FA deviation maps (overlaid on T2 FLAIR images) of an example of an FA outlier subject (left column) and an example of a normal FA subject (right column). (**c**) Structural images showing radiological findings in representative WMLV, FA, or MD outlier subjects of multiple sclerosis (a WMLV, FA, and MD outlier), lacunar infarcts with moderate small vessel disease (a WMLV, FA, and MD outlier), cyst (an MD outlier), and encephalomalacia (a FA and MD outlier). (**d**) WMLV, FA, or MD outlier interesting for follow-up. Left column: T2 FLAIR images of an outlier subject with severe biparietal non-typical distributed white matter lesions of uncertain etiology (a WMLV, FA, and MD outlier). Right column: regional FA deviation map (overlaid on T2 FLAIR images) of an FA outlier subject that was radiologically normal. FA was anomalously low in the genu of corpus callosum and the cause was unknown. For the regional FA deviation maps in (b) and (d), each map visualizes how the FA values in a subject deviate from the autoencoder-predicted FA values. For display purposes, in FA deviation maps, each white matter ROI is displayed in its full size instead of only the TBSS skeleton.

A proportion of the white matter outliers without any data acquisition or processing errors were reviewed by the neuroradiologist, and most of them were identified with radiological findings: This includes all of the reviewed WMLV outliers, 94.6% (35/37) of the reviewed FA outliers, and all of the reviewed MD outliers (Table 2). For instance, likely multiple sclerosis was identified in a subject who was an outlier in WMLV, FA, and MD (Fig. 5c). The diagnosis of multiple sclerosis was confirmed by the UKB health outcomes data. Lacunar infarcts and moderate small vessel disease were identified in another subject who was also an outlier in all three white matter-based imaging phenotypes (Fig. 5c). A parahippocampal cyst was identified in an MD outlier subject (Fig. 5c). Encephalomalacia (Fig. 5c) was identified in a subject who was an outlier in both FA and MD.

The etiology of the findings in some white matter outliers was uncertain. For example, the left panel of Fig. 5d shows an outlier subject in WMLV, FA, and MD measures, who was read as having severe biparietal atypically distributed white matter lesions of unknown etiology. Other than the subjects with radiological findings, a small number of the white matter outlier subjects reviewed appeared normal to the neuroradiologist (Fig. 5d, right panel, and Fig. S8b). For example, the right panel of Fig. 5d shows an FA outlier subject had an anomalously low FA value in the genu of corpus callosum specifically, but his T1w and T2-FLAIR images were normal-appearing. All these outliers of unknown etiology and normal-appearing outliers would be interesting for follow-up to determine the mechanism or whether they eventually present with specific clinical symptoms. Taken together, these results indicate that the non-artifactual outliers of white matter-based imaging phenotypes were associated with a large variety of different radiological findings. Normal-appearing outliers, each with unique FA or MD patterns, only constituted a small fraction of the white matter outliers.

### 3.6 Individual outliers of cortical thickness

We next examined the individuals with outlying CTh. As an example, Fig. 6a shows regional CTh deviation maps of an outlier subject versus a normal subject (about 4.9 * *IQR* away between these two subjects in CTh outlier score distribution). Widespread negative CTh deviations, representing thinner cortices in these regions, were observed in this outlier subject. The five CTh outlier subjects with good data collection/processing quality, were identified with prominent sulci or atrophy (Fig. 6b). Taken together, these results suggest that the very few non-artifactual CTh outliers were all associated with radiological findings.

**Fig. 6.**
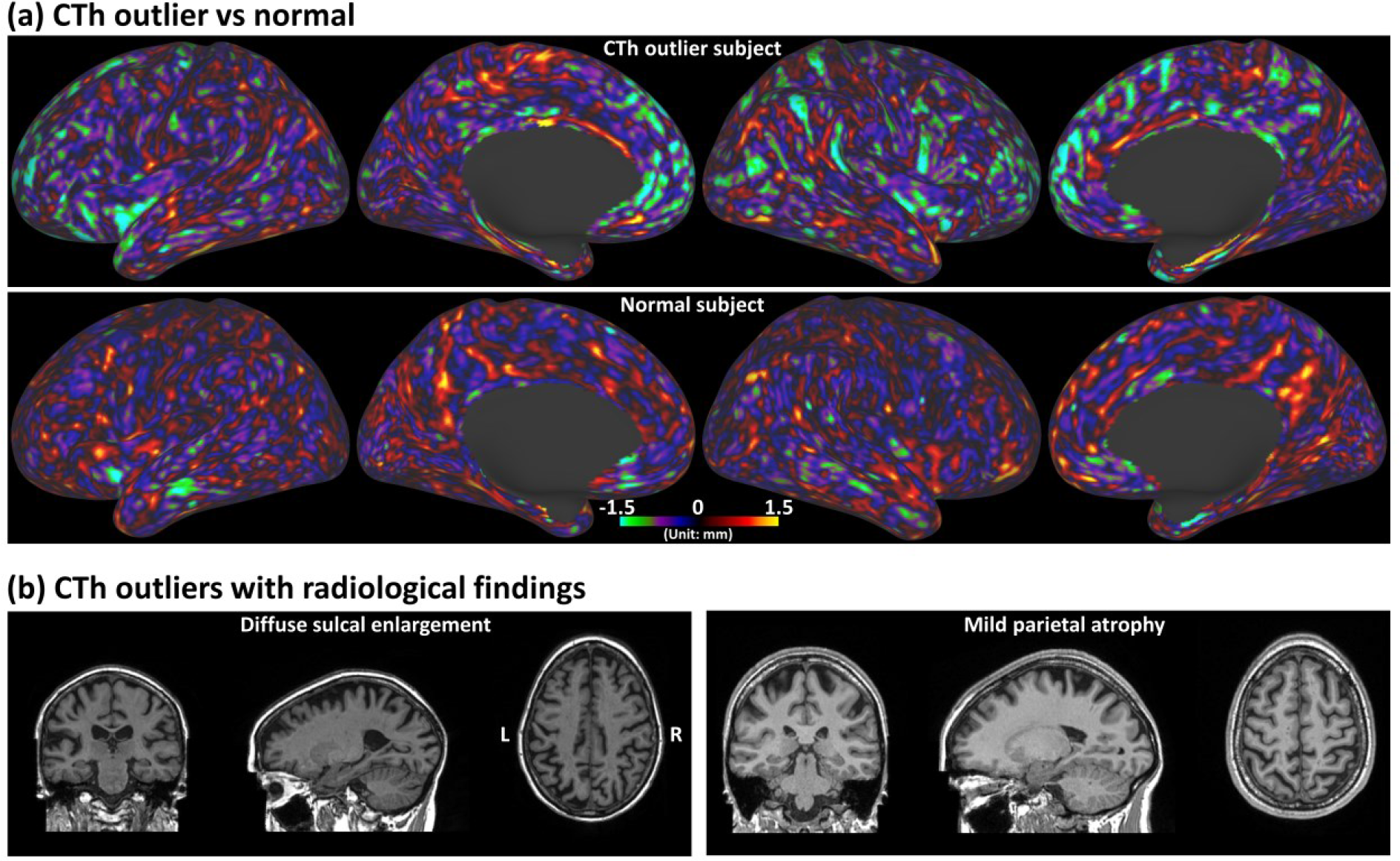
Individual outliers of CTh. (**a**) Regional CTh deviation maps (displayed on inflated cortical surfaces) of an example of a CTh outlier subject (first row) and an example of a normal CTh subject (second row). A regional CTh deviation map visualizes how the CTh values in a subject deviate from the autoencoder-predicted CTh values. (**b**) Structural images showing radiological findings in two representative CTh outlier subjects.

### 3.7 Outlier score relationships across imaging phenotypes

The relationship of outlier scores across different imaging phenotypes was assessed via pairwise Pearson correlation coefficients (Fig. 7a). Correlations between some white matter-based imaging phenotypes (FA versus MD; WMLV versus MD) were moderate (0.4 < r < 0.6), indicating they can capture similar outlying patterns in the white matter. All the other correlations were weak (0.2 < r < 0.4) or very weak (r < 0.2), indicating they were complementary and provided independent information.

**Fig. 7.**
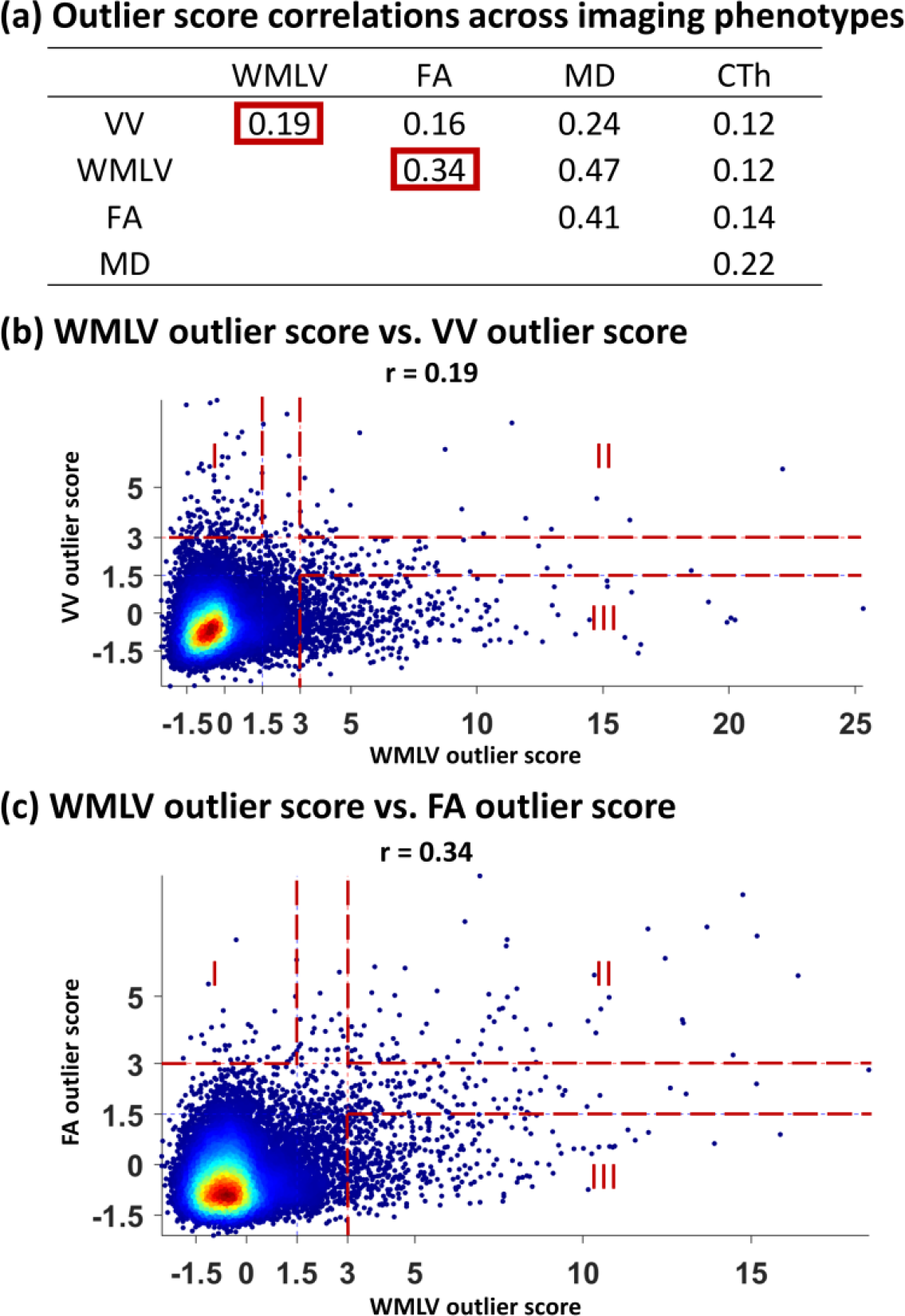
Relationship between outlier scores of different imaging phenotypes. (**a**) Correlations between the outlier scores of different imaging phenotypes. The outlier subjects with data collection/processing errors were not included in this analysis. The two representative relationships shown in (b) and (c) are encircled with red boxes. (**b**) WMLV outlier score plotted against VV outlier score. Zone I covered the VV outliers with normal WMLV. Zone II covered the subjects who were outliers in both VV and WMLV. Zone III covered the WMLV outliers with normal VV. (**c**) WMLV outlier score plotted against FA outlier score. Zone I covered the FA outliers with normal WMLV. Zone II covered the subjects who were outliers in both FA and WMLV. Zone III covered the WMLV outliers with normal FA.

To further illustrate these relationships, Fig. 7b shows a scatterplot of WMLV versus VV outlier scores, which were poorly correlated (r = 0.19). Very few subjects were both VV and WMLV outliers, as evidenced by the sparser data in Zone II than Zone I or III (Fig. 7b). Indeed, the density of Zone II was significantly lower than the other two zones (p ≈ 0, one-way analysis of variance [ANOVA] of 100000 bootstrap samples. Fig. S9a). It is therefore likely that the biological processes that led to large increases in WMLV are commonly independent of those that led to very enlarged VV. To illustrate another weak correlation, Fig. 7c shows a scatterplot of WMLV versus FA outlier scores (r = 0.34). The density of Zone II was significantly lower than Zone III (p ≈ 0, one-way ANOVA of 100000 bootstrap samples. Fig. S9b) but was close to Zone I. Fig. S9c shows two examples of the outliers in Zone II. The upper panel of Fig. S9c shows a subject that was an outlier in both VV and WMLV. This subject, diagnosed with ventriculomegaly and moderate white matter disease, had both periventricular and deep white matter lesions. The lower panel of Fig. S9c shows a subject that was an outlier in all VV, WMLV, FA, and MD. The radiological read determined there was small vessel disease evidenced by the white matter lesions, and possible Alzheimer’s disease evidenced by the parieto-temporal atrophy.

## 4. Discussion

In this study, a semi-automated, two-level outlier detection and screening methodology was used to investigate outliers in the MRI phenotypes of VV, WMLV, FA, MD, CTh, and RSFC (Fig. 1). Outlier score distributions were all more outlier-prone than a standard normal distribution (Fig. 2). Except for RSFC, outlier scores of these imaging phenotypes had good-to-excellent reliability as assessed by test-retest ICC of outlier scores (Fig. 3). Due to the low test-retest reliability of RSFC outlier scores, RSFC outliers were excluded from further individual-level analyses (Figs. 3 and S3). VV outliers had no data collection/processing errors and were all associated with radiological findings (Figs. 4 and S7). The white matter-based outliers were associated with more data collection/processing errors (Figs. S4 and S5) or radiological findings (Fig. 5c), though a small fraction of the white matter-based outliers appeared normal to the neuroradiologist (Figs. 5d, right panel, and S8b). CTh outliers were mostly due to data collection/processing errors (Fig. S6). The outlier scores of different imaging phenotypes were mostly independent, indicating that they each added information (Fig. 7).

### 4.1 Evaluation of unsupervised outlier detection

A common practice to evaluate an unsupervised outlier detection approach is to apply the method to a labeled dataset to calculate outlier scores without using the labels first, and these labels were used later as the ground truth when evaluating the performance of the unsupervised method (Aggarwal, 2017; Goldstein & Uchida, 2016). However, in the present study, the UKB data were unlabeled. Gibson et al. examined 1000 subjects of this cohort radiologically (Gibson et al., 2017), but unfortunately, we were unsuccessful in obtaining their radiological annotations. It should be noted that the outliers defined in the present study were composed of not only the subjects with radiological findings, but also the subjects with data collection/processing errors, as well as the radiologically normal-appearing outlier subjects, who still had large deviations from the group average (Fig. 5d, right panel). These radiologically normal-appearing outlier subjects are interesting because they could be the ones at higher risk to develop noticeable pathologies (de Groot et al., 2013). Therefore, instead of using any existing labels, we opted to evaluate our approach by quantifying how well the outlier scores align with the amounts of deviations from the group averages. For unidimensional imaging phenotypes, the outlier scores were linearly transformed from unidimensional volume measurements, so they exactly quantified such deviations. For multidimensional imaging phenotypes, the amount of individual deviations can be measured by converting the imaging phenotype into absolute z-scores and then averaging the absolute z-scores within each subject, and the autoencoder-derived outlier scores were found to be strongly correlated with such individual deviations (see Fig. S10 for details). The outlier scores also showed larger dynamic ranges than the amounts of deviations from the group averages (slopes > 1; Fig. S10), indicating that they were more sensitive in distinguishing small differences of outlierness than using the amounts of deviations from the group averages. Taken together, these results confirm that the outlier scores reliably characterized the degree of individual deviations from the group averages in this unlabeled dataset.

### 4.2 The approach to screen individual outliers in a large neuroimaging dataset

Large-scale neuroimaging datasets have emerged in recent years, with anywhere from 1,000 (Di Martino et al., 2014; Holmes et al., 2015; Van Essen et al., 2013) to more than 10,000 subjects (Hagler et al., 2019; Miller et al., 2016). Most studies using these datasets generally focus on the average imaging characteristics at a group level. There has been much less work on studying outlying individuals and the associated imaging phenotypes in these large neuroimaging datasets (Marquand et al., 2016; Mourao-Miranda et al., 2011; Pinaya et al., 2019; van Hespen et al., 2021). To begin to fill this gap, we set out to investigate individual outliers from about 20,000 UKB subjects.

Outlier detection was performed for all the commonly used, well-established brain imaging phenotypes. The data of most imaging phenotypes were well-curated, as evidenced by weak or very weak correlations between outlier scores and confounding factors (Fig. S11). The head motion and brain registration-related confounding factors evaluated here were similar to the ones described in recent work on confound modeling of the UKB brain imaging data (Alfaro-Almagro et al., 2021). However, there was inevitably a small fraction of data with acquisition or processing errors. Recent work has paid attention to the quality control of big neuroimaging datasets (Maximov et al., 2021), and it is critical to find ways to automatically identify the outliers associated with data collection/processing problems. In our work, this was achieved by screening the outliers via visual inspection and multiple different data quality control metrics (head motion level, brain registration quality, etc.; see *Materials and Methods* for details) to ensure the capture of different types of errors, including wrong FOV (Fig. S5a), head motion artifact (Figs. S4a, S5b, and S6a), incorrect segmentation (Figs. S4b and S6b), and incorrect registration (Figs. S5c and S6c). In addition, the outlier score assigned to each individual can be utilized as a useful summary index for assessing the effectiveness of different processing strategies at the individual level. For example, three common processing strategies regarding the global signal in RSFC were evaluated in this way (Figs. 3g-h and S3). Taken together, our method is valuable for curating large neuroimaging datasets.

A neuroradiologist read the structural images of outlier subjects that did not have data collection/processing errors. A large percentage (98.1%, 106/108) of the outlier subjects being read had radiological findings, such as large ventricles, masses, cysts, white matter lesions, infarcts, encephalomalacia, and prominent sulci. Most of these brain pathologies likely would have led to a recommendation to see a physician for follow-up. For example, a VV outlier subject (Fig. 4b) was diagnosed with a colloid cyst causing hydrocephalus, and the neuroradiologist’s read recommended this individual to see a neurosurgeon for follow-up. The aforementioned manual radiological screening study (Gibson et al., 2017) of the first 1000 UKB subjects showed that only 1.8% of the UKB subjects screened via systematic radiologist review had radiological findings in their brain MRIs. The much higher percentage (98.1%) of the subjects identified with radiological findings among our outlier subjects indicates that our method can effectively identify a subgroup that is greatly enriched with radiological findings from a large dataset.

### 4.3 Potential underlying mechanisms of outlier subjects with unknown etiology or were radiologically normal

Among the outliers identified with radiological findings, a few presented with unknown etiology. For example, eight UKB and five HCP VV outlier subjects had very large ventricles of uncertain etiology. The VV of these UKB subjects, ranging between 87.4 mL and 142.4 mL, was comparable to the upper range of VV in Alzheimer’s disease patients (Schott et al., 2005). The VV of these HCP subjects was between 45.4 mL and 56.2 mL, which were still much larger than the volumes of normal young healthy subjects.

Interestingly, the data also showed unexplained variations of VV between two monozygotic twin pairs in these HCP VV outlier subjects. Two female individuals within the first monozygotic twin pair had anomalously large VV (Fig. 4c, right panel), suggesting shared congenital, developmental, or environmental causes. In another monozygotic twin pair, only one twin had anomalous large VV (Fig. S7b). This is probably due to environmental influences or a de novo mutation early in development.

Another interesting case of unknown etiology was a UKB subject who was an outlier for WMLV, FA, and MD (Fig. 5d, left panel). In this subject, severe bilateral, confluent, and symmetrical white matter lesions were identified in the parietal white matter. Such lesion patterns were different from small vessel disease or multiple sclerosis, but were similar to reported cases of X-linked adrenoleukodystrophy (Geraldes et al., 2018). In the health outcomes data, this male subject was also reported to have hearing loss, a possible symptom of X-linked adrenoleukodystrophy, again indicating the possibility of this rare genetic disorder in this outlier subject with unknown etiology.

Two UKB FA outliers, two HCP FA outliers, and one HCP MD outlier were not identified with any data collection/processing issues or radiological findings, which are potentially interesting for investigating the underlying mechanisms of their large FA or MD deviations. For these FA outliers, anomalously low FA values were found either in the corpus callosum, superior longitudinal fasciculus, cingulum, posterior thalamic radiation, or limbs of the internal capsule (Figs. 5d, right panel, and S8b). A previous study showed that low FA in normal-appearing white matter preceded the conversion of low FA regions into white matter lesions (de Groot et al., 2013). Therefore, these FA outlier subjects may be at risk to develop lesions later in the regions of anomalously low FA. For the MD outlier subject, many small perivascular spaces were found on his structural MRI image. These perivascular spaces were not abnormal but could be responsible for the increased MD. Taken together, all the outliers discussed above would benefit from follow-up assessments to study underlying mechanisms and to see if they progress to any known clinical phenotype.

### 4.4 Generalizability of outlier detection to new UKB subjects

Since the UKB will ultimately enroll 100,000 subjects for brain imaging, it is important to verify that the outlier detection method used on the first 20,000 subjects in the present study can be applied to the rest of this population. Therefore, we made use of the second 20,000 UKB subjects released recently as a separate group for assessing the generalizability (referred to as *UKB replication group*; see Table S3 for detailed demographic information). These two groups were of comparable size and had no overlapping subjects. For each unidimensional imaging phenotype, generalizability was assessed by directly comparing the outlier score distribution obtained from the discovery group against the distribution obtained from the replication group, and no significant difference was found between these two distributions (Fig. S12a-b. Two-sample Kolmogorov-Smirnov tests: for VV, p = 0.36; for WMLV, p = 0.66). For each multidimensional imaging phenotype, first, the generalizability of the discovery group subjects’ outlier scores was evaluated by the ICC between two sets of their outlier scores calculated separately using two different autoencoders: one autoencoder trained using the discovery group itself, and another autoencoder trained using the replication group. The results showed that the ICC ranged from 0.86 to 0.99 (Fig. S12c-e). Second, the generalizability of the replication group subjects’ outlier scores was evaluated by the ICC between two sets of their outlier scores calculated separately using two different autoencoders: one autoencoder trained using the replication group itself, and another autoencoder trained using the discovery group. The results showed that the ICC ranged from 0.90 to 0.98 (Fig. S12c-e). Taken together, these results indicate that the UKB replication group is consistent with the UKB discovery group, and the results suggest that our trained outlier detection models can be generalized to new UKB subjects.

### 4.5 Technical considerations

There is a technical consideration about image preprocessing. For the subjects without usable T2-FLAIR images, ventricles were segmented using only T1w images. This choice should not affect the outlier detection of VV, because in our additional analysis of comparing T1w-only ventricle segmentation versus T1w and T2w combined ventricle segmentation in 18,328 UKB subjects, a close-to-one correlation (r = 0.999) was found between the VV values obtained from these two approaches.

### 4.6 Conclusions

The present study characterized individual outliers across multiple brain MRI phenotypes from 20,000 subjects. Every subject was parameterized with an outlier score per imaging phenotype to quantitate the outlierness. Outlier score distributions were all more outlier-prone than a standard normal distribution. The approach enabled the assessment of test-retest reliability via the outlier scores, which ranged from excellent reliability for VV, WMLV, and FA, to good reliability for MD and CTh. RSFC was excluded for individual-level outlier screening due to its low test-retest reliability. The individual-level analyses of the outliers revealed that a significant number of outliers were due to data collection/processing errors, demonstrating the usefulness of outlier detection in curating large neuroimaging datasets. Most of the remaining non-artifactual outliers were due to different brain pathologies as determined by a neuroradiologist, indicating that our approach can effectively identify a subgroup that is greatly enriched with radiological findings from a large unlabeled cohort. Several outliers had unknown etiology or were normal-appearing, and these outliers are candidates for follow-up to determine the mechanism or whether they eventually progress to a clinical phenotype. Our analysis suggests that unsupervised outlier detection of large neuroimaging datasets is valuable for data curation, reliability assessment, and identification of individuals for medical follow-up or further study of novel mechanisms. Outlier detection methods should contribute to the effort of developing automatic processes to analyze and interpret brain imaging data in large population cohorts.

## Acknowledgments

This study was supported by NIH/NINDS Intramural Research Program (Project numbers: NS002989 and NS003119). We thank Dr. Adam Thomas for helping with access to the UK Biobank datasets and for providing data storage resources. This research was conducted using the UK Biobank data under application number 22875. HCP data were provided [in part] by the HCP, WU-Minn Consortium (PIs: David Van Essen and Kamil Ugurbil; 1U54MH091657) funded by 16 NIH Institutes and Centers that support the NIH Blueprint for Neuroscience Research; and by the McDonnell Center for Systems Neuroscience at Washington University. In addition, the authors would like to thank UKB-Neuroimaging and HCP-Users mailing lists for helpful information. This work used NIH Biowulf high-performance computing resources (https://hpc.nih.gov).

## Abbreviations

VV: ventricular volume
WMLV: white matter lesion volume
FA: fractional anisotropy
MD: mean diffusivity
CTh: cortical thickness
RSFC: resting-state functional connectivity
UKB: UK Biobank
HCP: Human Connectome Project

