## Supplementary Information for "Outlier detection in multimodal MRI identifies rare individual phenotypes among 20,000 brains"

This file includes:

Supplementary Text: page 2 – 3

Figure S1 to S12: page 4 – 18

Table S1 to S4: page 19 – 22

References: page 23

### Supplementary Text

#### Outlier detection and screening in the HCP dataset

3T brain MRI data were obtained from the HCP 1200 Subjects Release (1113 subjects: 550 males and 656 females; age 22–37). If available, each HCP subject's T1w MPRAGE (N = 1113), T2w SPACE (N = 1094), and SE-EPI dMRI images (N = 1065) were used. The data were acquired on a 3T Siemens Connectome Skyra MRI scanner, and the detailed acquisition protocols can be found elsewhere (Glasser et al., 2013). The HCP project was approved by the Institutional Review Board of Washington University, and informed consent was obtained from all participants.

The following four commonly used brain imaging phenotypes were extracted from the HCP imaging preprocessing outputs: VV, FA, MD, and CTh. Because of the lack of T2-FLAIR in the HCP data, and poor WMLV segmentation accuracy when only using T1w images (Hotz et al., 2021), WMLV was excluded from the outlier detection of the HCP dataset. The detailed imaging phenotype extraction procedures are described as follows. The raw T1w MPRAGE and T2w SPACE images were preprocessed by the HCP structural pipeline (v4) (Glasser et al., 2013) based on FreeSurfer (v6) (Fischl, 2012). VV and CTh were extracted from the pipeline outputs. The dMRI data underwent FSL eddy-current and head-movement correction (Andersson & Sotiropoulos, 2016), gradient distortion correction, diffusion tensor model fitting (Basser, Mattiello, & LeBihan, 1994), and TBSS analyses. The skeletonized FA and MD images were averaged within the ROIs from the John Hopkins University white matter atlas (Mori et al., 2008). The FA or MD maps of 27 major white matter ROIs (Table S2) were extracted.

Using the similar primary screening procedures described in *Section 2.3* of the main text, outlier scores were calculated for each subject per imaging phenotype. Similar covariate control was performed, and this resulted in regressing out brain volume from VV. The total number of extreme outliers across all individual imaging phenotypes was 11. They were all distinct, because no subject was an outlier in more than one imaging phenotype. These outliers were checked individually using the similar secondary screening procedures described in *Section 2.5* of the main text, and none of them were associated with data collection/processing errors. The structural images of all these outlier subjects were reviewed radiologically, and the cases that would be interesting for follow-up were highlighted in Figs. 4c (the right panel), S7a (the first panel), S7b, and S8b (the second, third, and fourth panels). Table S4 summarizes the outlier detection and screening results in the HCP dataset.

### Supplementary Figures

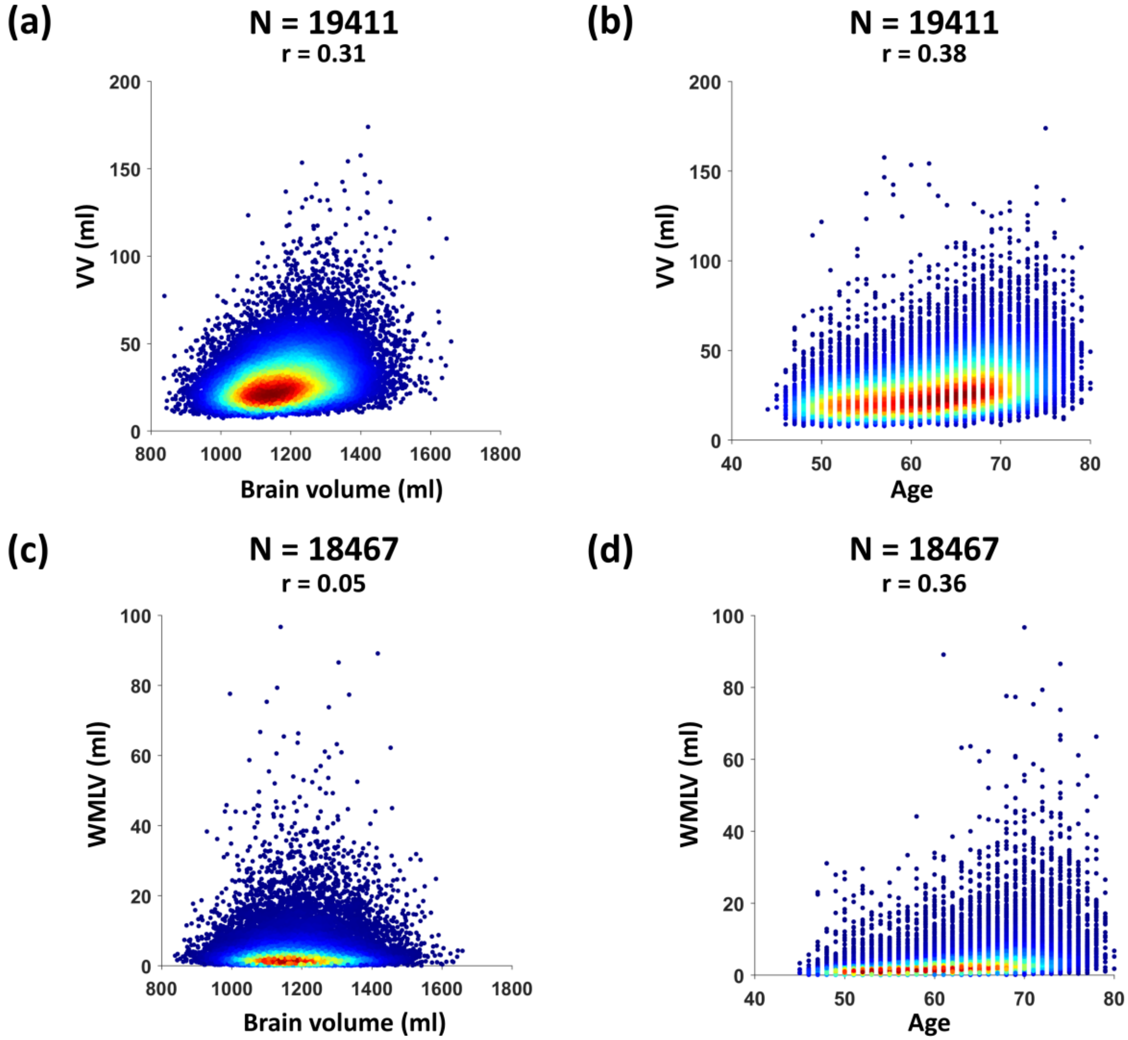


**Fig. S1.** The scatterplots between unidimensional imaging phenotypes (VV, WMLV; before covariate regression) and two covariates (brain volume, age). (**a**) Brain volume versus VV. (**b**) Age versus VV. (**c**) Brain volume versus WMLV. (**d**) Age versus WMLV.

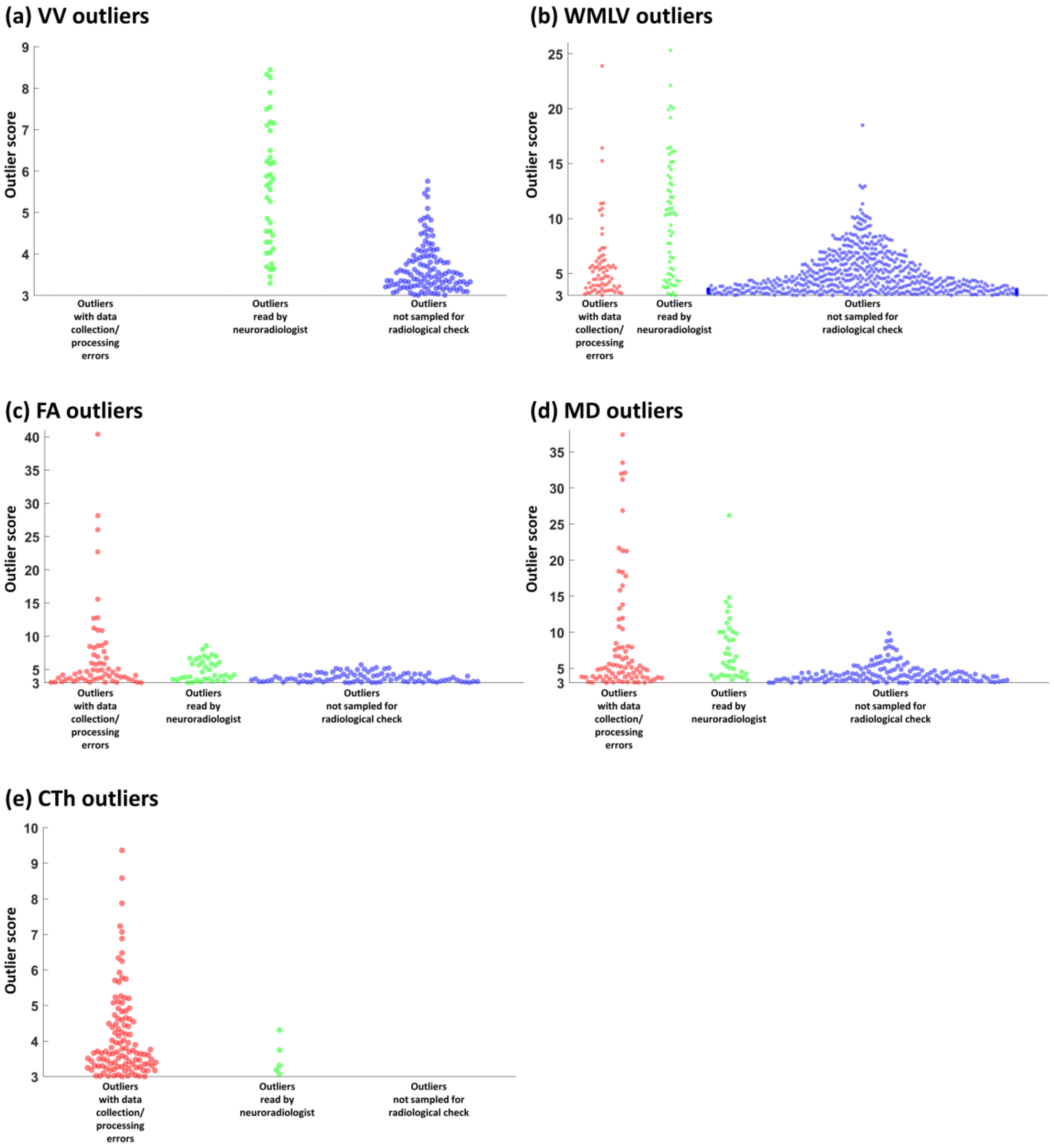


**Fig. S2.** Beeswarm plots for showing outlier subjects' outlier score ranges. The outliers with data collection/processing errors are represented by red dots. The non-artifactual outliers reviewed by a neuroradiologist are represented by green dots. The non-artifactual outliers not sampled for radiologically check are represented by blue dots. (**a**) VV outliers. (**b**) WMLV outliers. (**c**) FA outliers. (**d**) MD outliers. (**e**) CTh outliers.

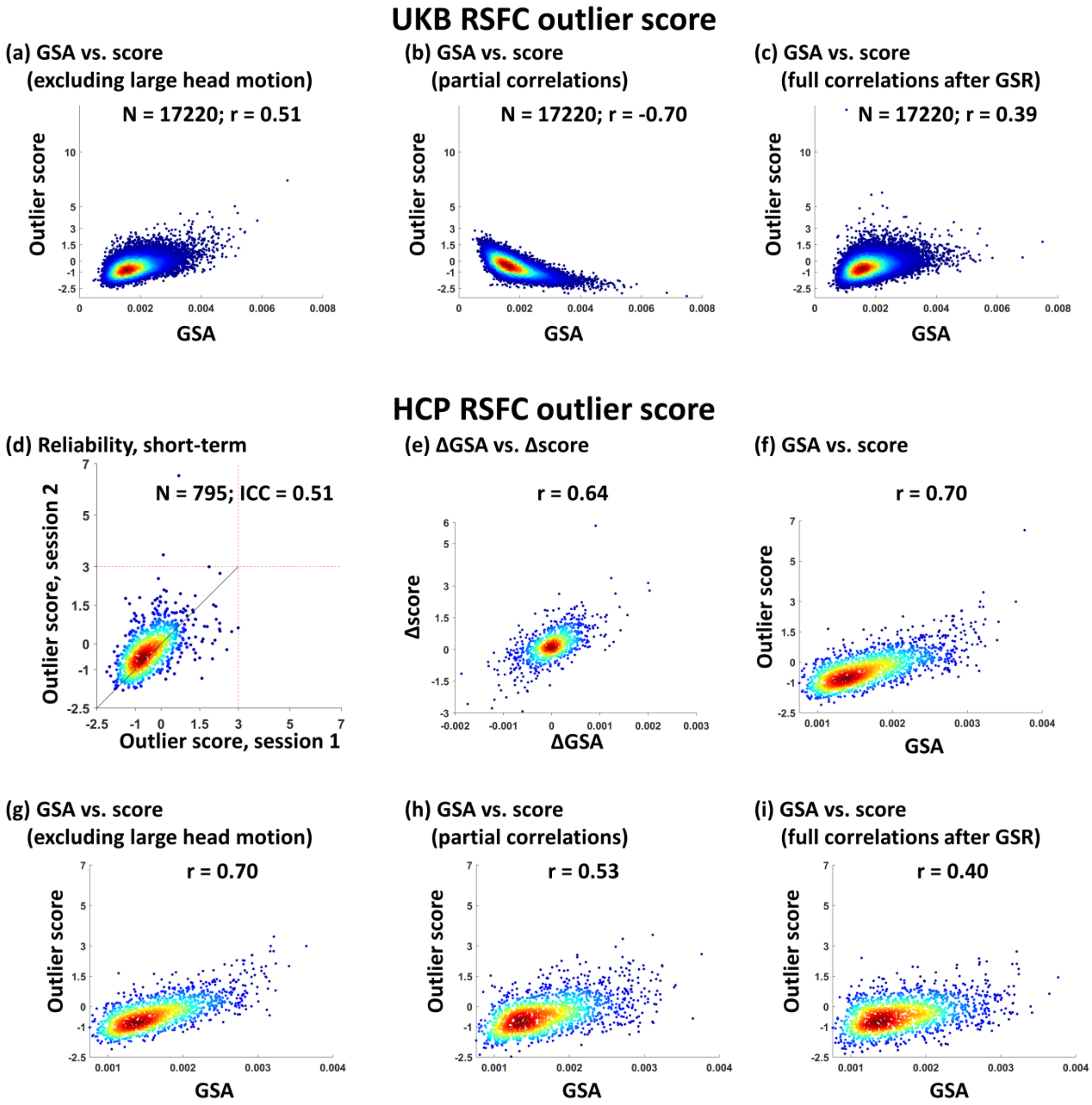


**Fig. S3.** Additional information on low test-retest reliability of RSFC outlier scores. (**a**) The scatterplot of global signal amplitude (GSA) versus RSFC outlier score in the UKB discovery group after excluding the subjects with large head motion (RSFC calculated using full correlations). (**b**) The scatterplot of GSA versus RSFC outlier score in the UKB discovery group (RSFC calculated using partial correlations). (**c**) The scatterplot of GSA versus RSFC outlier score in the UKB discovery group (RSFC calculated using full correlations after global signal regression [GSR]). Panels (d)-(i) show the assessment of test-retest reliability of RSFC outlier scores in the HCP dataset. The rsfMRI data of 795 HCP subjects using an improved image reconstruction algorithm "r227" were used. The data were preprocessed by the HCP functional pipeline (v3) (Glasser et al., 2013) and were denoised by ICA + FIX. Each HCP subject had two rsfMRI sessions, and the two runs within each session were demeaned, variance normalized, and concatenated temporally. Using the Gordon parcellation scheme (Gordon et al., 2016), RSFC was quantified by the Pearson cross-correlation coefficient with or without global signal regression, respectively. RSFC was also quantified using partial correlations with Tikhonov regularization (ρ = 0.01; FSLNets) (Pervaiz, Vidaurre, Woolrich, & Smith, 2020). The upper triangular parts of these RSFC matrices from each of the above three RSFC evaluation methods were extracted. The extracted RSFC data of the first sessions (aka "test") were used to train an autoencoder, and this trained autoencoder was applied to the data of the second sessions (aka "retest") to calculate retest outlier scores. (**d**) Test-retest reliability of RSFC outlier scores in the HCP dataset. Each subject's outlier score of the first session is plotted against the outlier score of the second session. Red dashed line: $Q3 + 3 * IQR$. (**e**) The scatterplot of test-retest GSA change versus test-retest RSFC outlier score change in the HCP dataset. (**f**) The scatterplot of GSA versus RSFC outlier score in the HCP dataset (RSFC calculated using full correlations). (**g**) The scatterplot of GSA versus RSFC outlier score in the HCP dataset after excluding the sessions with large head motion (RSFC calculated using full correlations). (**h**) The scatterplot of GSA versus RSFC outlier score in the HCP dataset (RSFC calculated using partial correlations). (**i**) The scatterplot of GSA versus RSFC outlier score (RSFC calculated using full correlations after GSR).

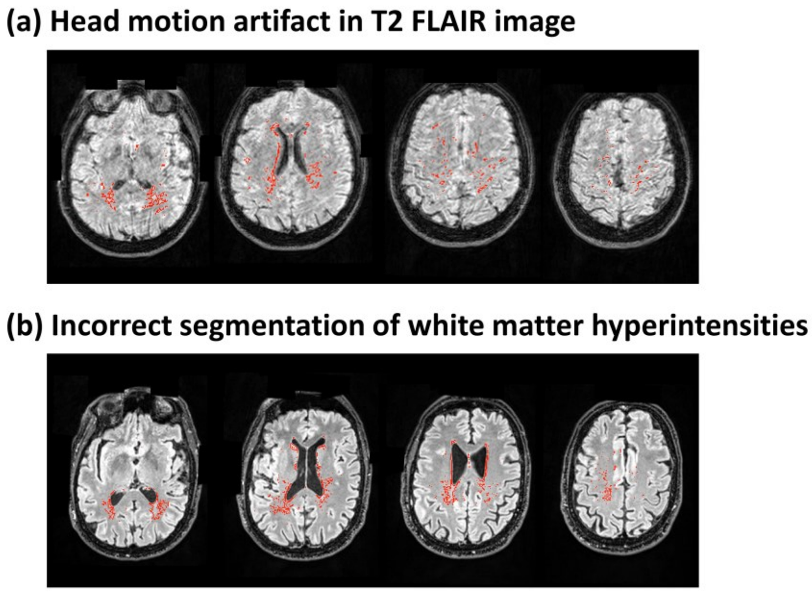


**Fig. S4.** Representative WMLV outliers associated with data collection/processing errors. The red line represents the inaccurate white matter lesion boundary segmented using BIANCA. (**a**) The segmentation of white matter hyperintensities in T2 FLAIR images was corrupted by head motion artifact. (**b**) Incorrect segmentation of white matter hyperintensities in T2 FLAIR images.

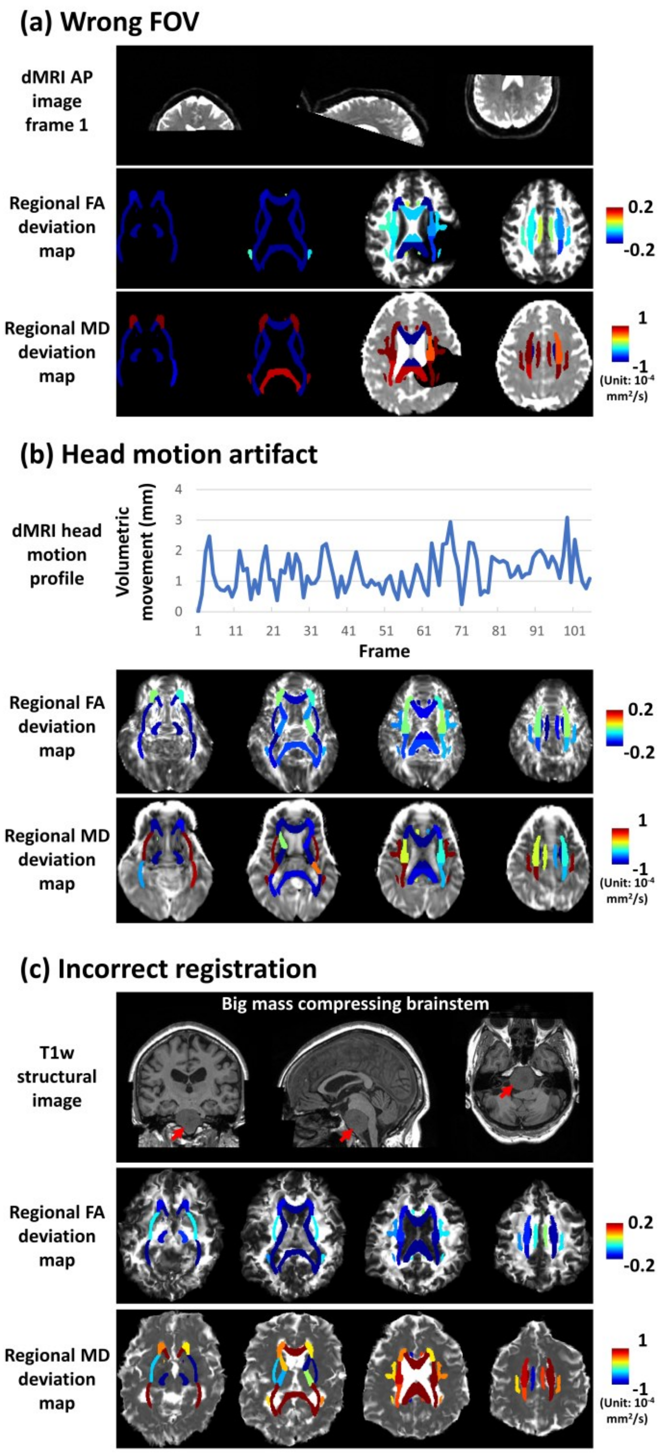


**Fig. S5.** Representative FA and MD outliers associated with data collection/processing errors. (**a**) Wrong FOV. Missing the inferior part of the brain in dMRI images (first row) resulted in anomalously large negative FA deviations (second row) and anomalously large positive MD deviations (third row) in these missing regions. (**b**) Head motion artifact. Large volumetric movements between adjacent dMRI frames (first row) corrupted the FA image (background image in the second row) and MD image (background image in the third row). This resulted in widespread anomalously large negative FA deviations (foreground map in the second row) and anomalously large positive MD deviations (foreground map in the third row). (**c**) Incorrect nonlinear registration. The registration was affected by a big mass pushing on the brainstem (first row). The FA and MD images after the registration were severely distorted (background images in the second and third rows) and this resulted in widespread anomalously large negative FA deviations (foreground map in the second row) and anomalously large positive MD deviations (foreground map in the third row). For display purposes, in these deviation maps, each white matter ROI is displayed in its full size instead of only the TBSS skeleton.

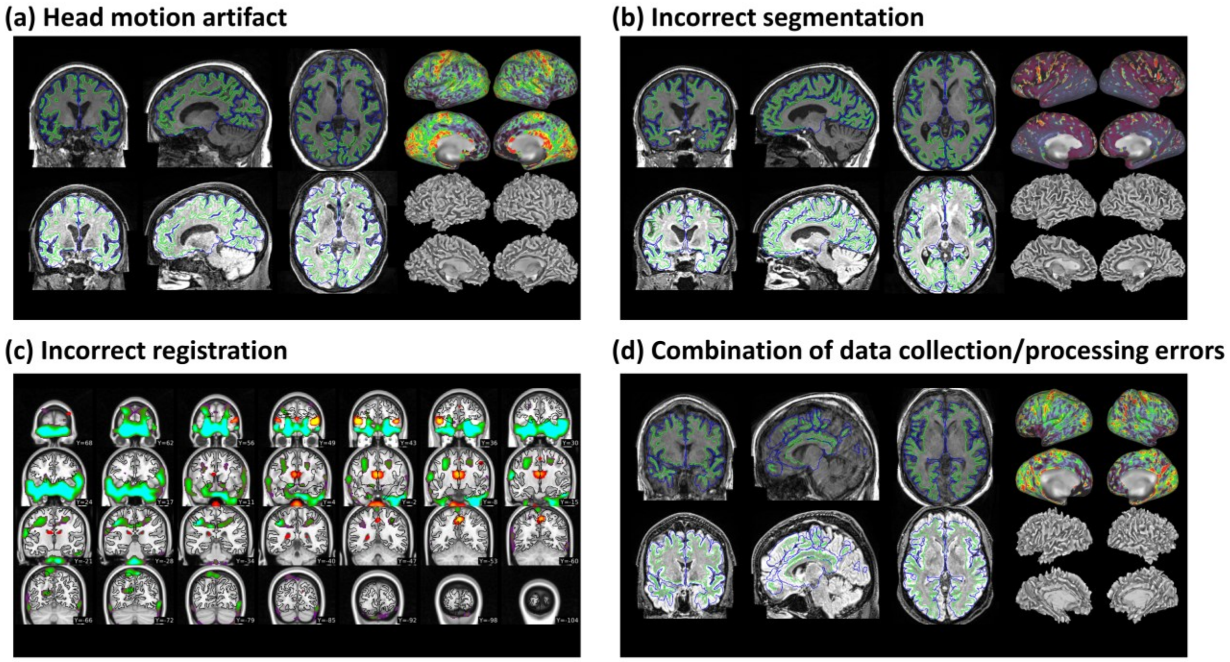


**Fig. S6.** Representative CTh outliers associated with data collection/processing errors. These figures are in the format of the HCP pipeline structural quality control scenes (https://github.com/Washington-University/StructuralQC; v1.4.0). (**a**) A CTh outlier subject with head motion artifact in structural images. (**b**) A CTh outlier subject with incorrect segmentation. (**c**) A CTh outlier subject with incorrect nonlinear registration of excessive volumetric deformation. (**d**) Combination of data collection/processing errors in a CTh outlier subject.

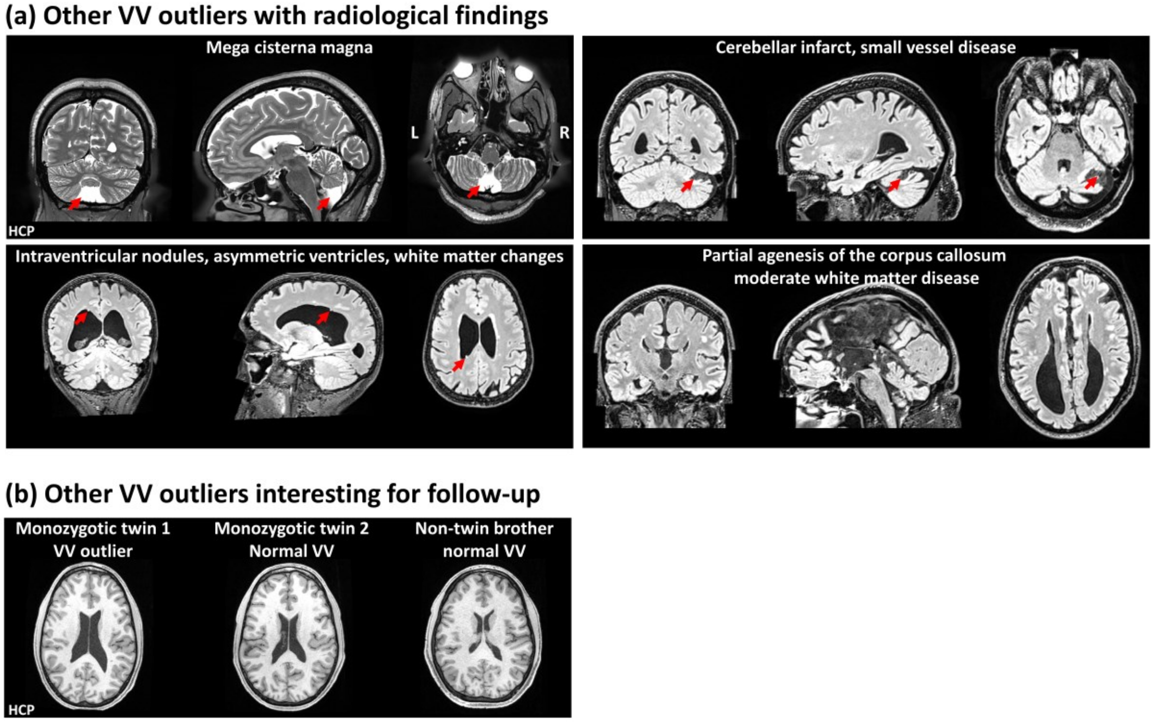


**Fig. S7.** Additional examples for VV outliers. (**a**) Structural images showing radiological findings in VV outlier subjects of a mega cisterna magna, an infarct, intraventricular nodules, and partial agenesis of the corpus callosum. The subject shown in the first panel is from the HCP dataset. (**b**) Structural images of other VV outlier subjects interesting for follow-up. These three subjects are from a family in the HCP dataset. One twin had large ventricles of unknown etiology, but the other twin and their non-twin brother had normal VV.

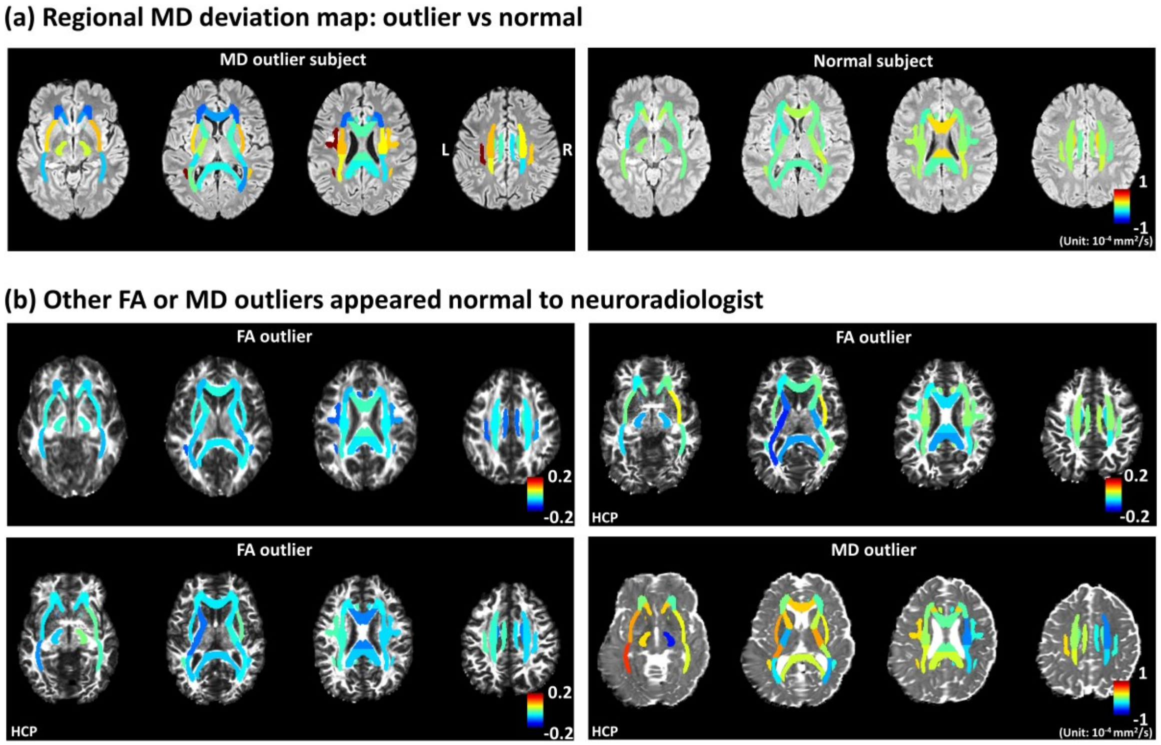


**Fig. S8.** Additional information on outlier detection using white matter-based imaging phenotypes. (**a**) Regional MD deviation maps (overlaid on T2 FLAIR images) of an example of an MD outlier subject (left column) and an example of a normal MD subject (right column). An MD deviation map visualizes how the MD values in a subject deviate from the autoencoder-predicted MD values. (**b**) Regional deviation maps (overlaid on FA or MD images) of other FA or MD outliers appeared normal to the neuroradiologist. The subjects shown in the second, third, and fourth panels are from the HCP dataset.

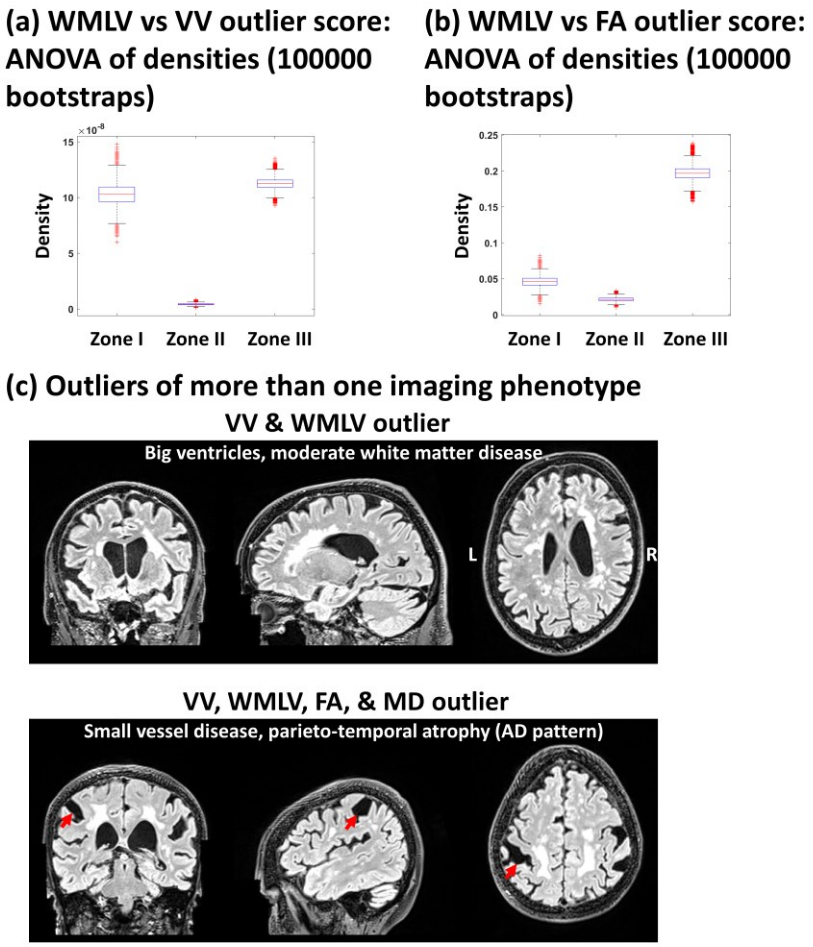


**Fig. S9.** Additional information on the relationship between outlier scores of different imaging phenotypes. (**a**) Bootstrapping results for comparing the subject densities in Zone I, II, and III in Fig. 7b. (**b**) Bootstrapping results for comparing the subject densities in Zone I, II, and III in Fig. 7c. (**c**) Structural images showing radiological findings in two subjects who were outliers in more than one imaging phenotype.

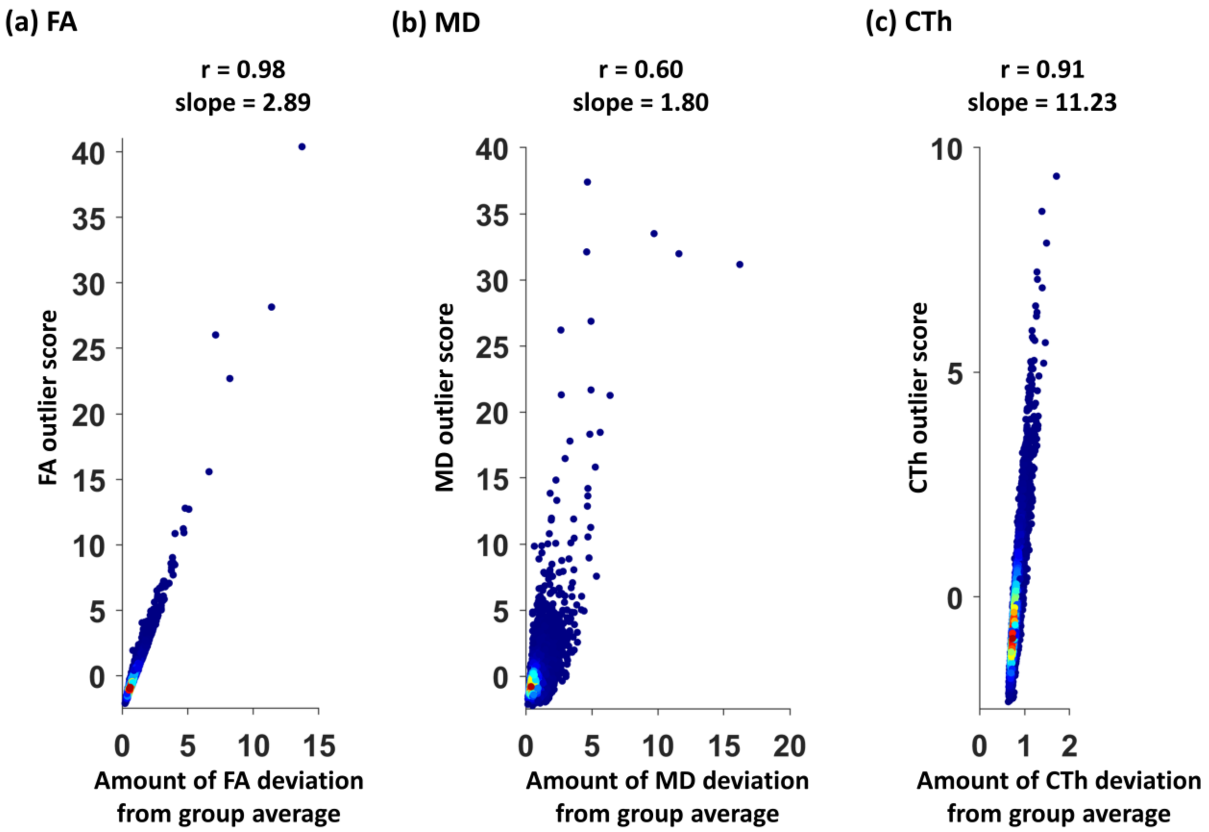


**Fig. S10.** Autoencoder-derived outlier scores were strongly correlated with the amounts of deviations from the group averages. For the calculation of the amount of deviations, using FA as an example, the FA value of each white matter ROI was converted to absolute z-score, and these absolute z-scores were averaged across ROIs within each subject. (**a**) The scatterplot of FA outlier score versus the amount of FA deviation from group average. (**b**) The scatterplot of MD outlier score versus the amount of MD deviation from group average. (**c**) The scatterplot of CTh outlier score versus the amount of CTh deviation from group average.

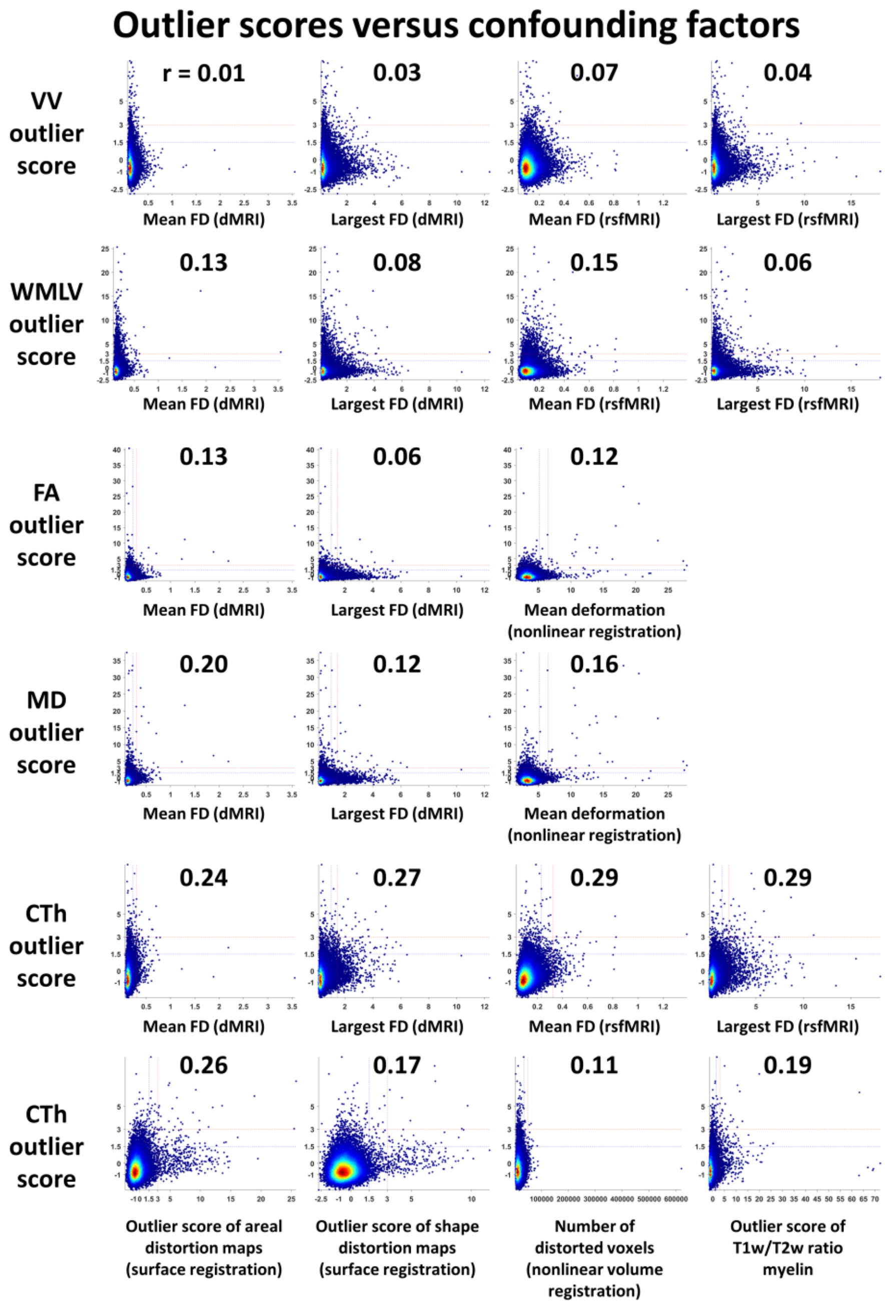


**Fig. S11.** Outlier scores versus confounding factors. Each small panel shows a scatterplot between the outlier score of an imaging phenotype (vertical axis) versus a confounding factor (horizontal axis), and the Pearson correlation between the two quantities is shown above each scatterplot. FD: framewise displacement (unit: mm).

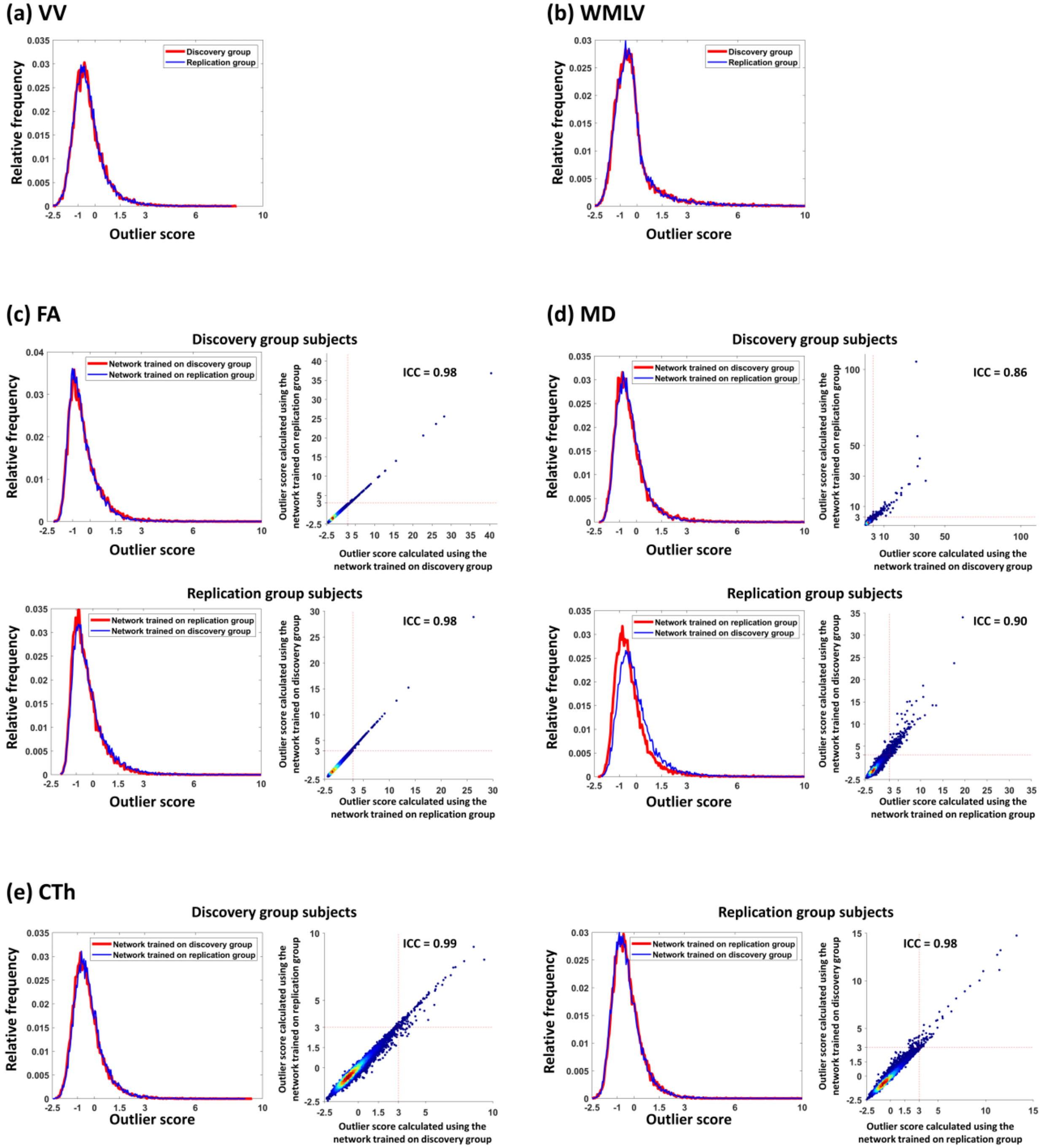


**Fig. S12.** Generalizability of outlier detection to new UKB subjects. (**a**) The distribution fitting of VV outlier scores of the UKB discovery group (red curve) overlaid on the distribution fitting of VV outlier scores of the UKB replication group (blue curve). (**b**) The distribution fitting of WMLV outlier scores of the UKB discovery group (red curve) overlaid on the distribution fitting of WMLV outlier scores of the UKB replication group (blue curve). (**c**) The UKB discovery group subjects' FA outlier scores (first row): The left panel shows the distribution fitting of the outlier scores calculated using the autoencoder trained on the discovery group itself (red curve) overlaid on the distribution fitting of the outlier scores calculated using the autoencoder trained on the replication group (blue curve). The right panel scatterplot shows these two sets of outlier scores plotted against each other. The UKB replication group subjects' FA outlier scores (second row): The left panel shows the distribution fitting of the outlier scores calculated using the autoencoder trained on the replication group itself (red curve) overlaid on the distribution fitting of the outlier scores calculated using the autoencoder trained on the discovery group (blue curve). The right panel scatterplot shows these two sets of outlier scores plotted against each other. (**d**) The UKB discovery group subjects' MD outlier scores (first row): The left panel shows the distribution fitting of the outlier scores calculated using the autoencoder trained on the discovery group itself (red curve) overlaid on the distribution fitting of the outlier scores calculated using the autoencoder trained on the replication group (blue curve). The right panel scatterplot shows these two sets of outlier scores plotted against each other. The UKB replication group subjects' MD outlier scores (second row): The left panel shows the distribution fitting of the outlier scores calculated using the autoencoder trained on the replication group itself (red curve) overlaid on the distribution fitting of the outlier scores calculated using the autoencoder trained on the discovery group (blue curve). The right panel scatterplot shows these two sets of outlier scores plotted against each other. (**e**) The UKB discovery group subjects' CTh outlier scores (first two panels): The first panel shows the distribution fitting of the outlier scores calculated using the autoencoder trained on the discovery group itself (red curve) overlaid on the distribution fitting of the outlier scores calculated using the autoencoder trained on the replication group (blue curve). The scatterplot in the second panel shows these two sets of outlier scores plotted against each other. The UKB replication group subjects' CTh outlier scores (last two panels): The third panel shows the distribution fitting of the outlier scores calculated using the autoencoder trained on the replication group itself (red curve) overlaid on the distribution fitting of the outlier scores calculated using the autoencoder trained on the discovery group (blue curve). The scatterplot in the fourth panel shows these two sets of outlier scores plotted against each other.

### Supplementary Tables

**Table S1.** Summary of the demographic information of the main dataset (UKB discovery group).

| Type | Number of subjects | Gender | | Age | | | | | |
|---|---|---|---|---|---|---|---|---|---|
| | | M | F | 40-49 | 50-59 | 60-69 | 70-79 | 80-89 | mean ± SD |
| T1w | 19411 | 9172 | 10239 | 782 | 6046 | 8791 | 3789 | 3 | 62.5 ± 7.5 |
| T2w-FLAIR | 18467 | 8701 | 9766 | 716 | 5767 | 8375 | 3606 | 3 | 62.5 ± 7.4 |
| dMRI | 17947 | 8430 | 9517 | 694 | 5631 | 8134 | 3485 | 3 | 62.5 ± 7.4 |
| rsfMRI | 17221 | 8074 | 9147 | 662 | 5412 | 7789 | 3355 | 3 | 62.5 ± 7.4 |

**Table S2.** Twenty-seven white matter ROIs from the John Hopkins University white matter atlas (Mori et al., 2008) were used in the outlier detection of FA and MD.

| ROI name |
|---|
| Genu of corpus callosum |
| Body of corpus callosum |
| Splenium of corpus callosum |
| Right cerebral peduncle |
| Left cerebral peduncle |
| Right anterior limb of internal capsule |
| Left anterior limb of internal capsule |
| Right posterior limb of internal capsule |
| Left posterior limb of internal capsule |
| Right retrolenticular part of internal capsule |
| Left retrolenticular part of internal capsule |
| Right anterior corona radiata |
| Left anterior corona radiata |
| Right superior corona radiata |
| Left superior corona radiata |
| Right posterior corona radiata |
| Left posterior corona radiata |
| Right posterior thalamic radiation (include optic radiation) |
| Left posterior thalamic radiation (include optic radiation) |
| Right sagittal stratum (include inferior longitidinal fasciculus and inferior fronto-occipital fasciculus) |
| Left sagittal stratum (include inferior longitidinal fasciculus and inferior fronto-occipital fasciculus) |
| Right external capsule |
| Left external capsule |
| Right cingulum (cingulate gyrus) |
| Left cingulum (cingulate gyrus) |
| Right superior longitudinal fasciculus |
| Left superior longitudinal fasciculus |

**Table S3.** Summary of the demographic information of the UKB replication group used in this study.

| Number of subjects | Gender | | Age | | | | | |
|---|---|---|---|---|---|---|---|---|
| | M | F | 40-49 | 50-59 | 60-69 | 70-79 | 80-89 | mean ± SD |
| 19350 | 9005 | 10345 | 104 | 5164 | 8164 | 5798 | 120 | 64.7 ± 7.4 |

**Table S4.** Summary of outlier detection and screening in the HCP dataset.

| Phenotype | | VV | FA | MD | CTh |
|---|---|---|---|---|---|
| Number of Subjects | | 1113 | 1065 | 1065 | 1094 |
| Skewness | | 1.50 | 0.92 | 0.68 | 0.33 |
| Kurtosis | | 6.64 | 4.42 | 3.71 | 3.14 |
| Number of outliers | | 8 (0.7%) | 2 (0.2%) | 1 (0.1%) | 0 (0%) |
| Outliers w/o data issue | | 8 | 2 | 1 | |
| Outliers read by neuroradiologist | | 8 | 2 | 1 | |
| Radiological comments | Large ventricles | 8 | | | |
| | White matter lesions | | | | |
| | Mass | | | | |
| | Cyst | 1 | | | |
| | Infarct | | | | |
| | Encephalo-malacia | | | | |
| | Prominent sulci | | | | |
| | Other findings | 2 | | | |
| | Normal | | 2 | 1 | |

Note: Empty entries are zeros.
